# Multivariate Bayesian Inversion for Classification and Regression

**DOI:** 10.1101/2025.05.09.653015

**Authors:** Joram Soch, Carsten Allefeld

## Abstract

We propose the statistical modelling approach to supervised learning (i.e. predicting labels from features) as an alternative to algorithmic machine learning (ML). The approach is demonstrated by employing a multivariate general linear model (MGLM) describing the effects of labels on features, possibly accounting for covariates of no interest, in combination with prior distributions on the model parameters. ML “training” is translated into estimating the MGLM parameters via Bayesian inference and ML “testing” or application is translated into Bayesian model comparison – a reciprocal relationship we refer to as *multivariate Bayesian inversion* (MBI). We devise MBI algorithms for the standard cases of supervised learning, discrete classification and continuous regression, derive novel classification rules and regression predictions, and use practical examples (simulated and real data) to illustrate benefits of the statistical modelling approach: interpretability, incorporation of prior knowledge, probabilistic predictions. We close by discussing further advantages, disadvantages and the future potential of MBI.

## 1 Introduction

The core aims of machine learning (ML) are the identification of patterns in data and the use of these patterns for the prediction of variables of interest [1], allowing for the automatization of complex problems such as optical character recognition [2], human speech recognition [3] or strategic game play [4]. Typically, ML is based on constructing a mapping from measured variables, or *features*, to some variables of interest, or *labels*. For example, ML may proceed by separating data points in a high-dimensional feature space to classify them into discretely labelled categories, as in support-vector machines (SVMs) [5, 6], or by transforming features using complex non-linear mappings to predict continuous target labels, as in deep convolutional neural networks (DNNs/CNNs) [7, 8].

In the ML approach, labels are algorithmically extracted from features. From a statistical perspective, an alternative is to model the features in terms of the labels (see Table 1) – and in many contexts where ML is applied, this is the more plausible approach, because the statistical model is often interpretable in terms of an underlying data-generating process. A statistical model entails the specification of probability distributions for the feature variables given the label variables, a so-called *generative model* [9, 10]. This typically invokes unknown parameters that can be estimated from the data and, in combination with prior distributions on those parameters, it becomes a *full probability model* [11]. Once the model parameters have been estimated, the model can be inverted for probabilistic inference of labels from features [12, 13]. Fundamental advantages of the statistical over the algorithmic approach are the possibility to incorporate explicit knowledge about the relationships between labels and features [14], both through the model structure and prior distributions, and its ability to qualify predictions by associating them with probabilities. For further technical advantages, see below and Section 5.

**Table 1.** Terminology. Analogous terms commonly used in the ML and statistics literatures, accompanied by an explanation of our use of them.

| Machine Learning | Statistics | Explanation |
| --- | --- | --- |
| features | dependent variables | measured variables which are the input to a predictive algorithm in machine learning or the output of a generative model in statistics |
| labels | independent variables | controlled (sometimes observed) variables which are thought to have some relationship with the features |
| classes | groups, categories | discrete labels |
| targets | covariate, predictor | continuous labels |
| feature matrix | data matrix | an observation-by-variable matrix containing dependent variables |
| label information | design matrix | an observation-by-regressor matrix containing independent variables |
| class labels | categorical design matrix | discrete categories represented by indicator regressors taking the values 0 or 1 |
| target values | parametric design matrix | continuous values represented by regressors taking real numbers as values |

The aim of this paper is to demonstrate this statistical approach by adapting a standard model from multivariate statistics to a predictive modelling context, showing its application in simulations and analyses of existing data sets. Multivariate features are assumed to be normally distributed and influenced by some label variable of interest as well as possibly covariates of no interest (see Table 1). Specifically, we use a multivariate general linear model (MGLM) [15, 16] to describe linear relationships between features, labels and covariates (see Table 2). Via a fully Bayesian treatment of the MGLM, we calculate conditional probabilities for the label variables, given the multivariate features. We refer to this as *multivariate Bayesian inversion* (MBI) and show how it can be used for both classification and regression.

**Table 2.** Notation. Symbols used in the mathematical description of MBI. Quantities are either measured (empirically observed), known (experimentally controlled), unknown (statistically estimated) or predicted (reconstructed from the features).

| Symbol | Dimension | Category | Meaning |
| --- | --- | --- | --- |
| $n, v, p$ | scalar | known | number of observations, variables and regressors |
| $Y$ | $n \times v$ | measured | data matrix |
| $X$ | $n \times p$ | known | design matrix |
| $B$ | $p \times v$ | unknown | regression coefficients |
| $E$ | $n \times v$ | unknown | error matrix |
| $V$ | $n \times n$ | known | observation covariance matrix |
| $\Sigma$ | $v \times v$ | unknown | feature covariance matrix |
| $P$ | $n \times n$ | known | inverse of $V$ |
| $T$ | $v \times v$ | unknown | inverse of $\Sigma$ |
| $x$ | $n \times 1$ | predicted | classes or targets |
| $Y_1, X_1$ | $n_1 \times \bullet$ | see above | $Y$ and $X$ in the training data |
| $Y_2, X_2$ | $n_2 \times \bullet$ | see above | $Y$ and $X$ in the test data |
| $y_{2i}$ | $1 \times v$ | measured | a single row of $Y$ in the test data |
| $x_{2i}$ | $1 \times p$ | predicted | a single row of $X$ in the test data |

In the process, we provide a systematic review of Bayesian parameter estimation and Bayesian model comparison for the MGLM. While these techniques are very well known [12, 17, 18] and have been applied in many different contexts before (e.g. [19, 20]), we here combine them to develop a comprehensive Bayes classifier (and Bayes regressor) for the MGLM.

Taken together, the contributions of this work are three-fold:

1. We rewrite the ML task of predicting from multivariate continuous data in terms of an interpretable probabilistic model which, rather than extracting labels from features, describes the effects that labels have on features.
2. We show how inversion of this model by means of Bayesian model selection can be used to perform ML tasks such as classification and regression and, in addition, provides posterior probabilities for the predicted labels.
3. We familiarize users of standard ML methods (e.g. SVMs) with statistical-model-based predictive analyses and demonstrate how common problems in ML – e.g. inclusion of variables of no interest, integration of prior knowledge, classification analyses for unbalanced categories, regression analyses for bounded variables, and correlated observations – can be handled naturally and elegantly by the MBI framework.

We are aware of the fact that the simple model which we describe here lacks several capabilities of currently leading ML methods (i.e. DNNs) – e.g. facilitating predicition in high-dimensional spaces, coping with non-normal and heteroskedastic error distributions, or allowing for non-linear relationships between features and labels via kernel-based methods. Our goal is not to outperform these approaches in terms of predictive accuracy. Rather, we want to provide a basic framework for statistical-model-based Bayesian machine learning which may be further extended to address these issues. Some possible extensions are discussed in Section 5.3.

The present paper is structured as follows. First, we outline the statistical theory and derive the mathematical equations underlying MBI (see Section 2). Next, we present a series of MBI applications based on the analysis of simulated and empirical data (see Sections 3 and 4). In these sections, we also compare results based on MBI with results obtained using SVMs, representing a comparably flexible and currently popular ML approach [1, 21, 22]. Finally, we review various aspects illustrated by these simulations and discuss advantages, disadvantages and possible extensions of MBI (see Section 5).

### Terminology

Note that, following the statistical modelling approach, we conceptualize an ML problem as learning a relationship of the form *Y* = *f* (*X*) + *ε* – in which *Y* are features and *X* are labels –, as opposed to the typical ML approach of extracting estimates of labels from observations of features, i.e. 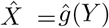. Consequently, we refer to labels as “independent variables” and features as “dependent variables”, not the other way around. This is because the statistical model describes *Y* as a function of *X*, but when inverted, can be used to deduce *X* from *Y*.

### Notation

In the following, we denote *n*-dimensional zero vectors as 0_*n*_ and *n*-dimensional vectors of all ones as 1_*n*_. Similarly, we denote *n*×*m* zero matrices as 0_*nm*_ and *n* × *m* matrices of all ones as 1_*nm*_. We write the *n*×*n* identity matrix as *I*_*n*_ and write its *i*-th column, i.e. the *i*-th *n*-dimensional unit vector, as *e*_*i*_ for *i* = 1, …, *n*. We use 𝒩 (*µ*, Σ) to denote the multivariate normal distribution with expected value *µ* ∈ ℝ^*n*^ and covariance matrix 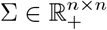 where 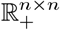 is the set of all positive-definite symmetric matrices of size *n* × *n*. Finally, we use 𝒩 ℳ (*M, U, V*) to denote the matrix normal distribution with expected value *M* ∈ ℝ^*n*×*m*^, covariance matrix across rows 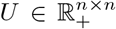 and covariance matrix across columns 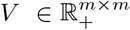.

## 2 Theory

In this section, we introduce the foundations of multivariate Bayesian inversion (MBI). We first review the notions of predictive algorithms and generative models and then introduce the multivariate general linear model as a generative model (see Section 2.1). We go on to describe how, using Bayesian data analysis, one can either estimate the parameters of such a model or perform comparison between the models themselves (see Section 2.2). Using the MBI framework for predictive analyses rests on the reciprocal estimation of parameters and labels, i.e. either label values are known and model parameters have to be estimated or model parameters are known and label values have to be predicted (see Section 2.3). Finally, upon partitioning available data into training and test data, MBI can be used for discrete classification or continuous regression, respectively (see Section 2.4).

### 2.1 Linear models for multivariate data

#### 2.1.1 Predictive algorithms vs. generative models

Consider some features *Y* ∈ 𝒴 and some labels *X* ∈ 𝒳, i.e. random variables (dependent and independent variables, see Table 1) with sample spaces 𝒴 and 𝒳, respectively. Then, a function *g*: 𝒴 → 𝒳 is called a *predictive algorithm*. In supervised learning, one typically calibrates *g* based on a training set of features and labels (*Y*_1_, *X*_1_) and then evaluates the performance of *g* by predicting the labels *X*_2_ of test features *Y*_2_,

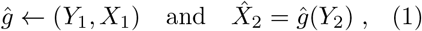

where *ĝ* denotes the calibrated predictive algorithm and 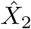 are the predicted labels of the test features [23, 24]. A common example for this is the *multivariate continuous prediction task* in which *Y* ∈ ℝ^*v*^ is a random vector of *v* continuous features and either *X* ∈ *{*1, …, *C*} is a discrete class index or *X* ∈ ℝ is a real number. In the former case, the multivariate continuous prediction task is referred to as a *classification task* and is typically solved using algorithms such as multinomial logistic regression or support vector classification [5, 25]. In the latter case, the multivariate continuous prediction task is referred to as a *regression problem* and can be solved using e.g. multiple linear regression or support vector regression [6].

Alternatively, the relationship between features *Y* and labels *X* can be furnished by taking a statistical perspective. Here, a statement about the probability distribution of *Y*, given *X* and some model parameters *θ* ∈ Θ, is referred to as a *generative model*. The term implies that, given fixed parameters *θ*, features *Y* can be generated by sampling the data-generating mechanism,

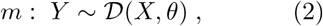

where *m* denotes the generative model and 𝒟 (*X, θ*) signifies an arbitrary probability distribution parametrized in terms of *X* and *θ*. This distribution implies a probability function *p*(*Y* |*X, θ, m*): 𝒴 → ℝ_≥0_, which – when viewed as a function *θ* for fixed *Y* – is referred to as the *likelihood function* of the generative model.

The idea behind this generative approach is to model the effects that labels have on features (*X→ Y*), with the intention to later invert the model in order to estimate labels from features (*Y →X*). When labels are latent variables causally influencing observable features – which is often, but not always the case in real-world applications (see Section 4) –, the generative model captures the data-generating mechanism by describing *p*(*Y*| *X*) rather than *p*(*X* |*Y*) (see Section 2.7).

A typical example for this is a Gaussian model where 𝒟 is a (potentially multivariate) Gaussian distribution and *θ* are weights or coefficients characterizing linear or non-linear effects of *X* on *Y*. Importantly, the fact that generative models are based on explicit probabilistic assumptions about features, labels and parameters makes them compatible to principled forms of model estimation and model selection such as maximum likelihood, Bayesian posterior inference, classical information criteria or Bayesian model comparison [11, 12, 18, 26, 27].

For example, in Bayesian inference, a *prior distribution* over model parameters *p*(*θ* |*m*) is specified which allows to infer the *posterior probabilities* of the model parameters, given the features *p*(*θ* |*Y, m*), or to calculate the *marginal likelihood* of the measured features, given the model *p*(*Y* |*m*).

In the following, we show how the multivariate general linear model can be written as a generative model and made amenable to Bayesian analyses, where the results of these analyses can then be employed as a means to solve multivariate continuous prediction tasks in the style of predictive algorithms.

#### 2.1.2 Multivariate general linear models

Let *Y* ∈ ℝ^*n*×*v*^ denote an observed random matrix of measured data comprising *n* observations of *v* features and let *X* ∈ ℝ^*n*×*p*^ denote a known matrix comprising *n* associated observations of *p* predictor variables (also called “regressors”) that possibly influence the measured data. Then, we define a multivariate general linear model (MGLM) *m* as

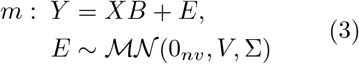

where *B*∈ ℝ^*p*×*v*^ denotes a matrix of unknown (regression) coefficients, *E* ∈ ℝ^*n*×*v*^ denotes and unknown feature an unobserved error matrix, and ℳ 𝒩 (0_*nv*_, *V*, Σ) denotes a matrix-normal distribution with zero mean, known observation covariance matrix 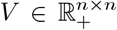 covariance matrix 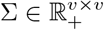.

In applications to data sets comprising multiple features for multiple experimental units (e.g. participants in a study), *V* models the error covariation between different experimental units and Σ models the error covariation between of features within each experimental unit. In applications to spatiotemporal data sets, where observations correspond to different time points and features correspond to different locations in space, *V* may be referred to as a *temporal covariance matrix* and Σ may be referred to as a *spatial covariance matrix* [16].

Note that, in equation (3), *X* and *V* are assumed to be known, whereas *B* and Σ are assumed to be unknown (see Table 2). This is compatible with the fact that observations, i.e. rows of *Y*, typically have some known autocorrelation structure or are assumed independent (hence, *V* is pre-specified), whereas features, i.e. columns of *Y*, typically have some unknown covariation which needs to be inferred (hence, Σ is estimated). The assumption of matrix-normally distributed errors also means that all observations have the same covariance across columns Σ and that all features have the same covariance across rows *V*. If observations are independent and identically distributed (i.i.d.), as is often assumed in ML applications, then *V* = *I*_*n*_ and the model simplifies to (see Appendix A)

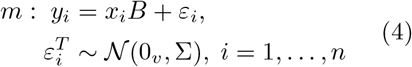

where *y*_*i*_ ∈ ℝ ^1×*v*^, *x*_*i*_ ∈ ℝ ^1×*p*^, *ε*_*i*_ ∈ ℝ ^1×*v*^ are row vectors and 𝒩 (0_*v*_, Σ) denotes a multivariate normal distribution with zero mean and covariance matrix Σ ∈ ℝ ^*n*×*n*^.

Since *E* follows a matrix-normal distribution and *XB* is constant, the MGLM formulated in (3) is equivalent to the following generative model (see Appendix A):

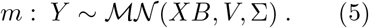

With the matrix-normal probability density function 𝒩 ℳ: ℝ ^*n*×*v*^ *→* ℝ_*>*0_, equation (5) then implies the following likelihood function for *m* (see Appendix A):

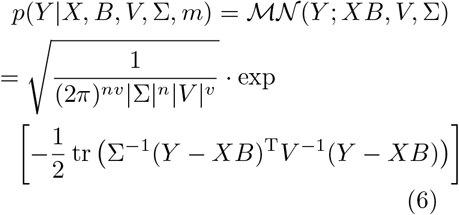

For reasons of mathematical convenience, we write this in terms of the precision matrices 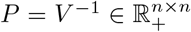 and 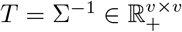 (see Table 2):

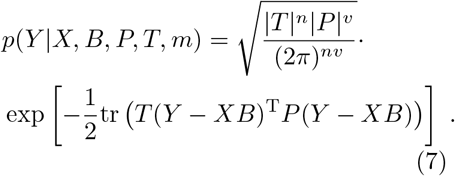

Note that, among the conditioned variables, *V* (or equivalently, *P*) and *m* are constant which is why, in the following sections, we will drop them for clarity of notation. In contrast to that, *B* and *T* are random entities described by some probability distribution. *X* is considered fixed when estimating *B* and *T* (see Section 2.3.1) and random when using a distribution over *B* and *T* to predict it (see Section 2.3.2). Taken together, we will therefore refer to the likelihood function of the MGLM as *p*(*Y* |*X, B, T*).

### 2.2 Bayesian inference for MGLMs

#### 2.2.1 Bayesian parameter estimation

In Bayesian inference, one usually specifies a prior density *p*(*θ* |*m*), i.e. a probability distribution for *θ* unconditional on *Y*, and the goal is to derive the posterior density *p*(*θ* |*Y, m*): Θ → ℝ_≥0_, i.e. the probability distribution for *θ* conditional on *Y*, given the model *m* [11, 18, 28–30]. Bayes’ theorem implies that this posterior density is proportional to the product of the likelihood function *p*(*Y θ, m*) and the prior density:

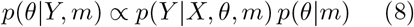

Following (7), the MGLM model parameters are 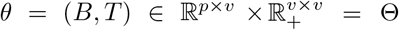 where 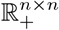 is the set of all positive-definite symmetric matrices of size *n* × *n*. Thus, Bayesian MGLM estimation requires a prior distribution on *B* and *T*. In order to obtain a tractable posterior, we will use a conjugate prior, i.e. a prior distribution that, when combined with the likelihood function, leads to a posterior distribution that belongs to the same family of probability distributions. For the MGLM, this conjugate prior is a normal-Wishart distribution on the coefficient matrix *B* and the precision matrix *T* ([11, ch. 3]; [31, sect. 8])

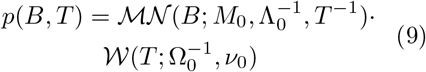

where *M*_0_ ∈ ℝ^*p*×*v*^ is the prior mean, 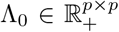 is the prior precision matrix, 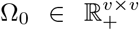 is the prior inverse scale matrix, *ν*_0_ ∈ ℝ are the prior degrees of freedom and 𝒩 ℳ: ℝ^*p*×*v*^ → ℝ_*>*0_ and 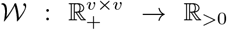 are the respective probability density functions.

Because this distribution is a conjugate prior, the application of (8) results in a posterior distribution that is also a normal-Wishart distribution (see Appendix B)

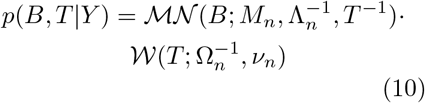

where the posterior parameters *M*_*n*_, Λ_*n*_, Ω_*n*_ and *ν*_*n*_ turn out to be functions of the prior parameters, the feature variables *Y* and the predictor variables *X* (see Appendix B):

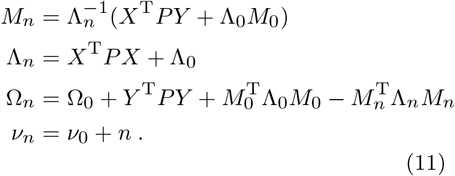

Note that, if observations are assumed to be i.i.d., we have 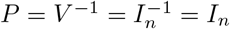 and the matrix *P* simply disappears from those equations. This is the case for most applications of linear regression models and will also be assumed throughout most parts of this paper (but see Simulation 4 in Section 3).

#### 2.2.2 Bayesian model comparison

In addition to obtaining posterior distributions for the model parameters *θ*, Bayesian inference also allows to perform statistical inference on the generative model *m* itself. This is achieved by calculating the marginal likelihood *p*(*Y* |*m*): 𝒴 → ℝ_≥0_, i.e. the probability of the data *Y*, independent of particular parameter values *θ*, given only the model *m* [12, 32–34]. The law of marginal probability implies that this marginal likelihood is an integral of the product of likelihood function and prior distribution:

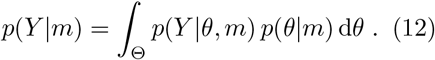

Application of (12) to the MGLM with 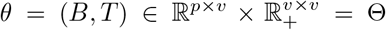 yields

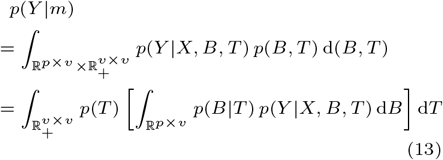

and the marginal likelihood with (11) becomes (see Appendix C)

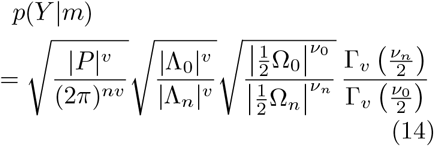

where Γ_*p*_: ℝ → ℝ is the multivariate Gamma function of order *p* ∈ ℕ. Once again, if observations are assumed i.i.d., such that rows of the data matrix *Y* are uncorrelated – as it is the case in most linear modelling contexts –, we have *P* = *I*_*n*_, such that the term |*P*| ^*v*^ in (14) would be replaced by 1.

If two or more models are differing by the predictor variables *X* they utilize (giving rise to different likelihood functions *p*(*Y* |*X, θ, m*)) or by the prior parameters (*M*_0_, Λ_0_, Ω_0_, *ν*_0_) they impose (giving rise to different prior distributions *p*(*θ* |*m*)), this consitutes a case for model comparison. Consider a model space ℳ = {*m*_1_, …, *m*_*M*_} where *M* is the number of models. Then, posterior model probabilities can be calculated from marginal likelihoods (14) using Bayes’ theorem

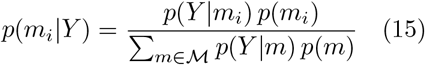

for *i* = 1, …, *M* where *p*(*m*_*i*_) are prior model probabilities, i.e. prior beliefs about the relative likelihoods of models *m* ∈ ℳ before seeing the data *Y*.

### 2.3 Multivariate Bayesian inversion

#### 2.3.1 Estimation of parameters, given the labels

In Section 2.2.1, we have introduced how Bayesian inference can be used for MGLM parameter estimation. In this section, we explain how this method constitutes the first step of a multivariate Bayesian inversion analysis.

The fact that, according to equation (7), the probability of the measured data *Y* is conditional on both the model’s regressors *X* and the model parameters *θ* = (*B, T*), means that both of these quantities can be considered either estimated and known or unknown and to-be-predicted. In some situation, e.g. in a designed experiment, it could be that regressor values are known and model parameters have to be learned. In other contexts, e.g. in consumer choice prediction, it could be that model parameters are known and regressor values have to be inferred.

Consider a data set (*Y, X, P*) ∈ ℝ^*n*×*v*^ ℝ^*n*×*p*^ ℝ^*n*×*n*^ specified by data matrix *Y*, design matrix *X* and precision matrix *P* (see Section 2.1.2). In this case, *X* would be considered known and *B* and *T* have to be estimated following (8):

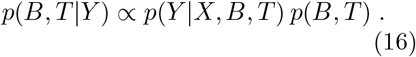

To do so, a prior density *p*(*B, T*) as in (9) has to be specified by fixing the prior parameters *M*_0_, Λ_0_, Ω_0_ and *ν*_0_ (see Section 2.2.1). If there is no prior knowledge, i.e. one does not have concrete prior beliefs about the model parameters *B* and *T*, one possibility is to use non-informative prior parameters [30] given by

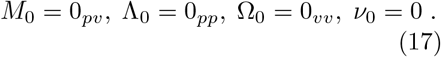

Using (10) and (11), the posterior distribution becomes

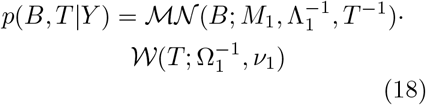

where the posterior parameters after seeing the data are (see Appendix D):

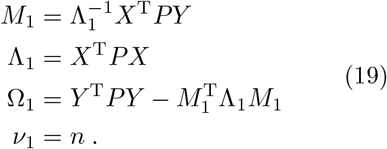

Note that the prior parameters (17) are specified in such a way that the posterior parameters (19) are only influenced by the observed data *Y* and the known regressors *X* in this example (cf. eq. 11). Moreover, as we only estimate one *T* = Σ^−1^, the feature covariance matrix Σ is assumed to be constant across observations, i.e. the same for all data points. For multivariate Bayesian classification (see Section 2.4.1), this means that MBI pools covariance across classes. For multivariate Bayesian regression (see Section 2.4.1), this means that the covariance is not dependent on the target variable.

#### 2.3.2 Identification of labels, given the parameters

In Section 2.2.2, we have introduced how Bayesian inference can be used for MGLM model comparison. In this section, we explain how this method constitutes the second step of a multivariate Bayesian inversion analysis.

Other than before, it could be that a distribution *p*(*B, T*) is available, for example (but not necessarily) because it has been estimated from independent data. In this case, the model parameters *B* and *T* can be assumed known – or at least partially known, or known with some uncertainty – when approaching new data.

Consider a single data point (*y*_*i*_, *x*_*i*_) ∈ ℝ^1×*v*^ × ℝ^1×*p*^ specified by a new observation *y*_*i*_ and the associated regressor values *x*_*i*_, where *i* is an index over previously unseen data. Let us assume that the goal is to identify the *x*_*i*_ belonging to a single new *y*_*i*_. For this one observation, the multivariate model reads (cf. eq. 4):

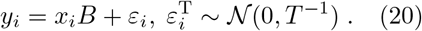

Now, *B* and *T* can be considered known in the sense that the distribution *p*(*B, T*) acts as the prior distribution for the new data point, such that the prior predictive distribution of the observed *y*_*i*_, given the unknown *x*_*i*_ follows from (12). For example, assume that the distribution *p*(*B, T*) is given by the posterior distribution *p*(*B, T* |*Y*) in (18) specified by the parameters *M*_1_, Λ_1_, Ω_1_ and *ν*_1_ from (19). Then, assuming this as our prior knowledge and using (14), the marginal likelihood *p*(*y*_*i*_|*x*_*i*_) becomes

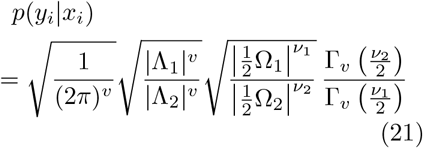

where the posterior parameters after seeing the new data are (see Appendix D):

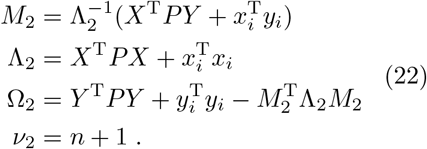

Crucially, different regressor values can be plugged in for *x*_*i*_ in equation (22), enabling to treat the identification of labels as a model selection problem. More precisely, we are testing competing models for the observations *y*_*i*_ by submitting different possibilities for the regressors *x*_*i*_, to see which of these possibilities maximizes the marginal likelihood *p*(*y*_*i*_|*x*_*i*_) and is thus the most likely value of *x*_*i*_ (for a similar approach, see [20]). Given that we can specify some prior knowledge about the predictor variables *x*_*i*_, the marginal likelihoods from (21) can be used to obtain posterior probabilities for label values *x*_*i*_, given the measured data *y*_*i*_:

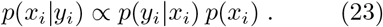

Note that, other than in equation (16), *X* is not considered fixed anymore, but *x*_*i*_ is now a random variable which has prior distribution *p*(*x*_*i*_) and can be estimated in the form of a posterior distribution *p*(*x*_*i*_|*y*_*i*_).

Multivariate Bayesian inversion (MBI) is the alternating procedure in which first, MGLM parameters are estimated, given known labels, and in which second, labels are infered on, given known MGLM parameters. In Sections 2.4.1 and 2.4.2, we will outline an application of this MBI framework for cross-validated predictive analyses which are based on splitting the data into training and test set in order to predict labels for left-out observations using either classification or regression.

### 2.4 Cross-validated MBI for prediction

#### 2.4.1 Multivariate Bayesian classification

Let 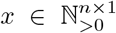 be a vector of non-zero class indices (1, 2, 3, …) corresponding to the observations in the data matrix *Y* ∈ ℝ^*n*×*v*^ (see Figure 1) and let 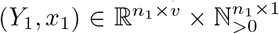 and 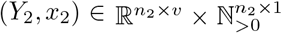 denote training and test set, respectively. The number of data points are *n*_1_ and *n*_2_, such that *n*_1_ + *n*_2_ = *n*.

**Fig. 1.**
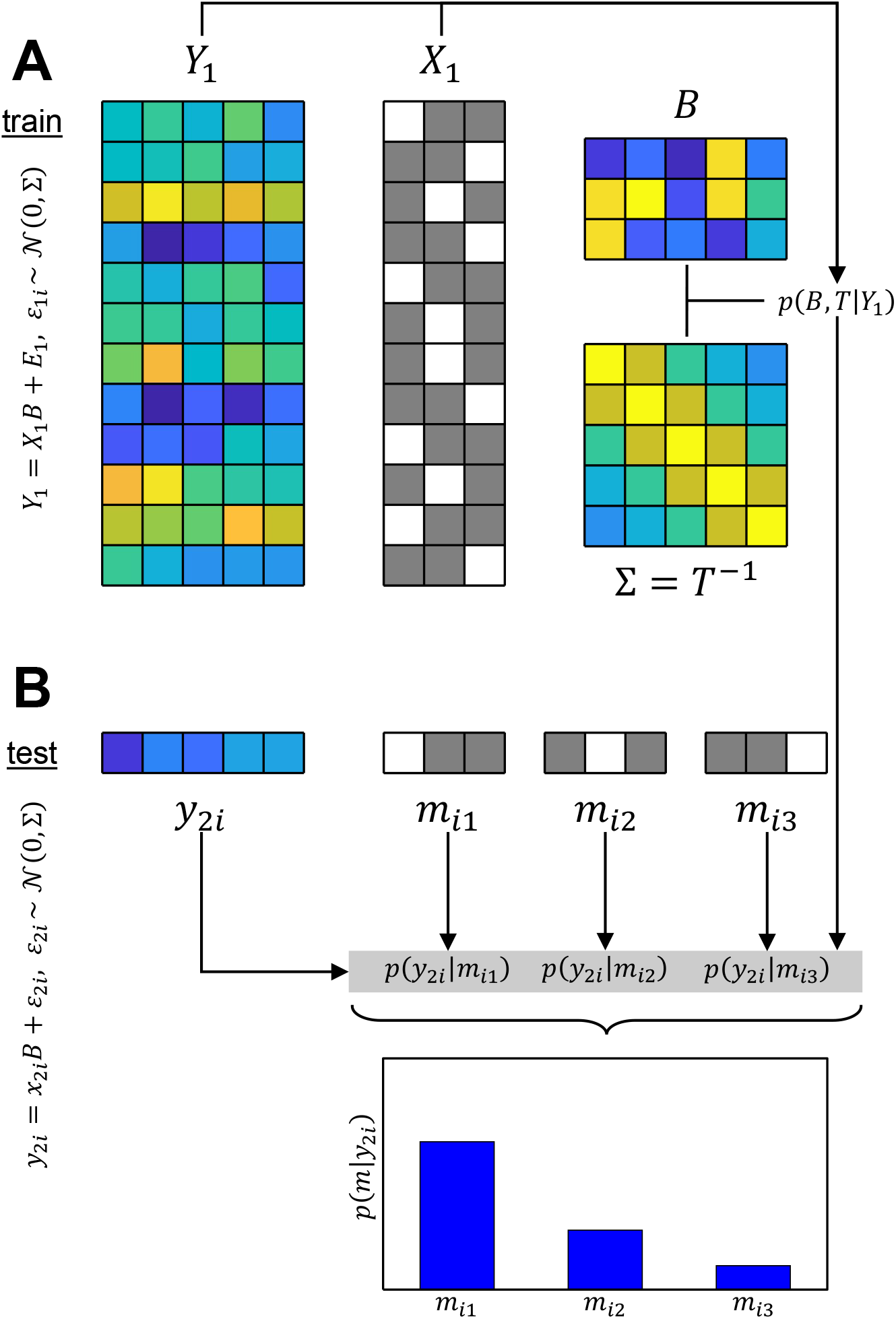
Multivariate Bayesian inversion for classification. **(A)** In the training set, a categorical design matrix *X*_1_ is formed and the posterior distribution over regression coefficients *B* and inverse feature covariance *T* = Σ^−1^ is calculated from the measured data *Y*_1_. **(B)** In the test set, each category *j* is successively tested against each data point *y*_2*i*_ as a possible model *m*_*ij*_, resulting in posterior class probabilities *p*(*x*_2*i*_ = *j* |*y*_2*i*_). Multivariate Bayesian classification is a two-step procedure, separately estimating parameters (top) and inferring on classes (bottom). Note that between-observation covariance, denoted in the text via *V* and *P*, has been omitted (i.e. set to *V* = *P* = *I*_*n*_) in this example for simplicity.

First, we define a “regressor generator function” 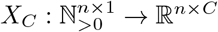 which sets the *j*-th column in the *i*-th row to 1, if the *i*-th observation belongs to the *j*-th class, where *C* = max(*x*) is the number of classes (see Figure 1A):

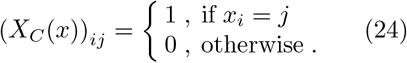

The result is a “categorical design matrix” in which each column vector is a binary indicator, i.e. only taking values 0 or 1, representing one class.

Since *x*_2_ is unknown, i.e. we do not know the classes in the test set, we have to specify a set of candidate models. Let *m*_*ij*_ be the model asserting that the single observation *y*_2*i*_ in the test set belongs to the *j*-th class (see Figure 1B):

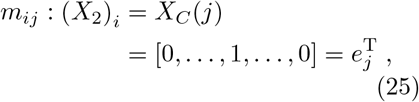

i.e. the *i*-th row of *X*_2_ is a 1 ×*C* vector of zeros with a one at the *j*-th position.

Then, we can calculate the marginal likelihood of the test data point *y*_2*i*_ based on (21):

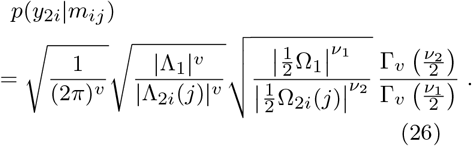

This allows to quantify posterior class probabilities for each observation based on (15):

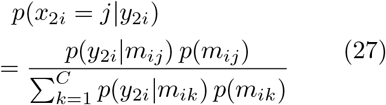

where *p*(*m*_*ij*_) is the prior probability that test set observation *i* belongs to class *j*.

Note that the only terms in (26) depending on the candidate model *m*_*ij*_ are

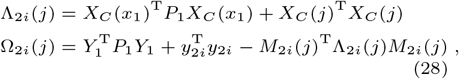

such that the posterior class probabilities from (27) can be written as

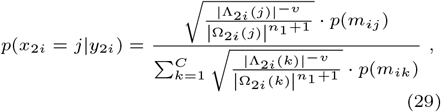

for all test data points *i* = 1, …, *n*_2_. If one were to predict the classes in the test set, picking the one which maximizes the posterior probability would be a reasonable choice:

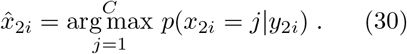

#### 2.4.2 Multivariate Bayesian regression

Let *x* ∈ ℝ^*n*×1^ be a vector of real-number target values (*x*_*i*_ ∈ ℝ, *i* = 1, …, *n*) corresponding to the observations in the data matrix *Y* ∈ ℝ^*n*×*v*^ (see Figure 2) and let 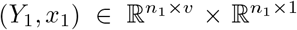 and 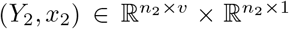 denote training and test set, respectively. The number of data points are *n*_1_ and *n*_2_, such that *n*_1_ + *n*_2_ = *n*.

**Fig. 2.**
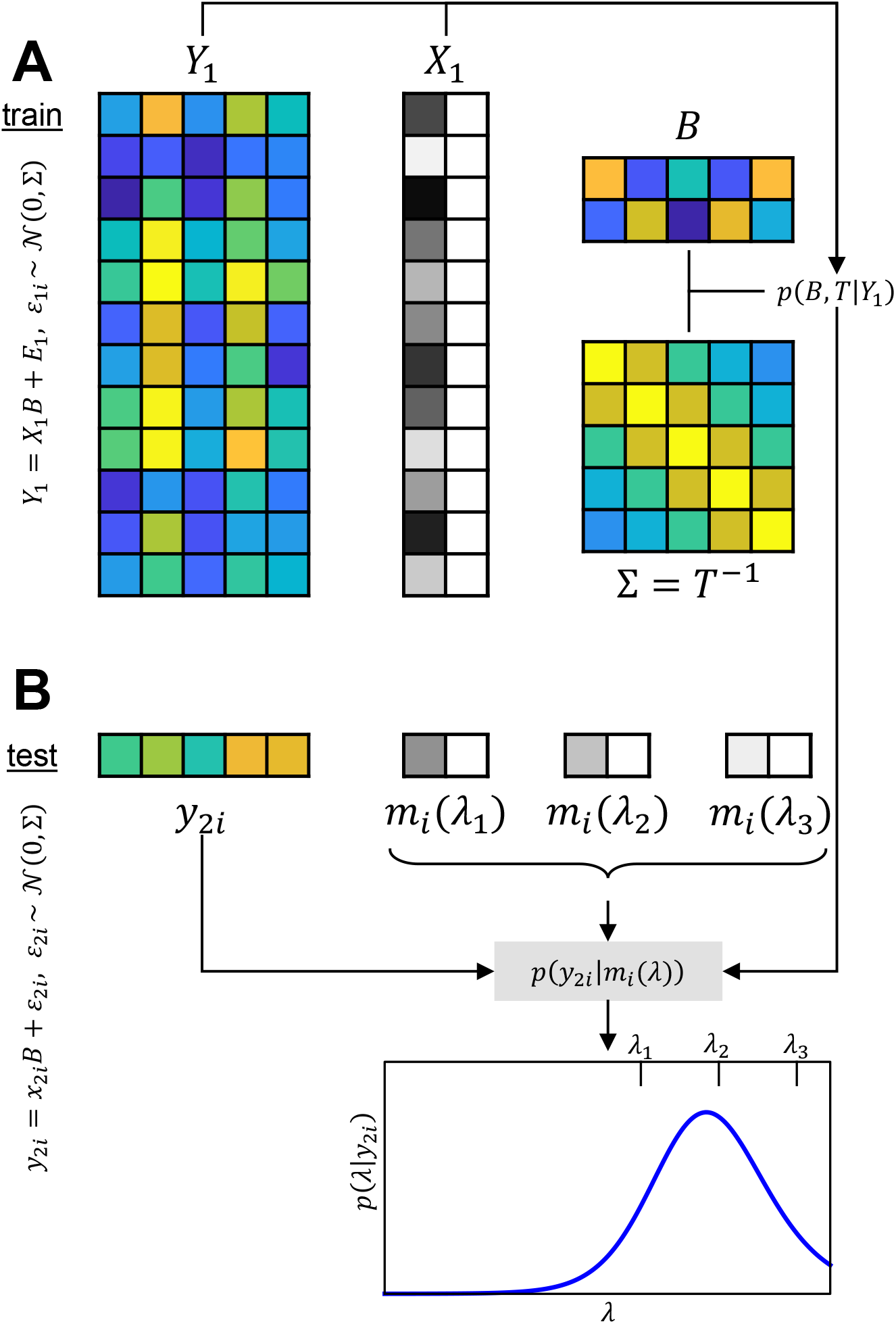
Multivariate Bayesian inversion for regression. **(A)** In the training set, a regression design matrix *X*_1_ is formed and the posterior distribution over regression coefficients *B* and inverse feature covariance *T* = Σ^−1^ is calculated from the measured data *Y*_1_. **(B)** In the test set, the marginal likelihood for each data point *y*_2*i*_ is a continuous function of the possible target values *λ*, resulting in a posterior target density *p*(*x*_2*i*_ = *λ*| *y*_2*i*_). Target values corresponding to the three models shown are marked on top of the posterior density plot. Multivariate Bayesian regression is a two-step procedure, separately estimating parameters (top) and inferring on targets (bottom). Note that between-observation covariance, denoted in the text via *V* and *P*, has been omitted (i.e. set to *V* = *P* = *I*_*n*_) in this example for simplicity.

First, we define a “regressor generator function” *X*_*R*_: ℝ^*n*×1 →^ ℝ^*n*×2^ which horizontally concatenates the target vector and a vector of ones (see Figure 2A):

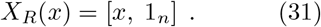

This results in a “regression design matrix” in which the first column represents the target variable and the second column is a constant regressor.

Since *x*_2_ is unknown, i.e. we do not know the targets in the test set, we have to specify a set of candidate models. Let *m*_*i*_(*λ*) be the model asserting that the target value belonging to the single observation *y*_2*i*_ is equal to *λ* (see Figure 2B):

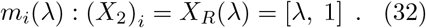

i.e. the *i*-th row of *X*_2_ is a 1× 2 vector with *λ* as first entry and one as second entry.

Then, the marginal likelihood of the test data point *y*_2*i*_ is a function of *λ* based on (21):

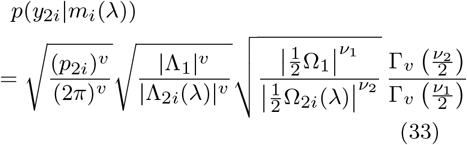

This allows to derive a posterior target density over *λ* for each observation based on (15):

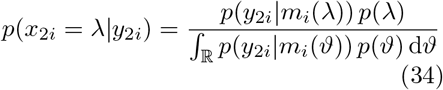

where *p*(*λ*) and *p*(*ϑ*) is the prior probability density over the *i*-th test set label *x*_2*i*_.

Note that the only terms in (33) depending on the candidate value *λ* are

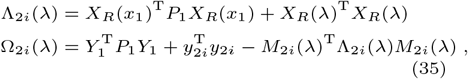

such that the posterior target density from (34) can be written as

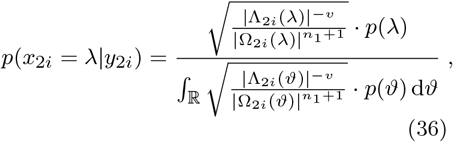

for all test data points *i* = 1, …, *n*_2_. If one was to predict the target value for each data point in the test set, reporting either the posterior expected value or the maximum-a-posteriori estimate would be reasonable choices:

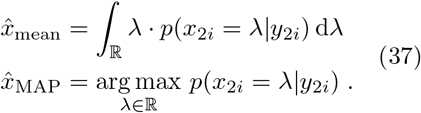

### 2.5 Inclusion of covariates

In Sections 2.4.1 and 2.4.2, MBC and MBR were formulated without covariates, i.e. with only class indices *x* or target values *x* influencing the measured data *Y*. We will now write down the design matrices for MBI with inclusion of covariates, i.e. incorporating confound variables *Z* which are not themselves the target of prediction, but may have influences on the available features *Y*.

Let *Z* ∈ ℝ^*n*×*c*^ be a matrix of covariates corresponding to the observations in the data matrix *Y* ∈ ℝ^*n*×*v*^ and let 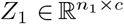 and 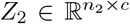 denote covariate values from training and test set, respectively. Let (*Z*_2_)_*i*_ be the 1 × *c* vector of covariate values for the *i*-th observation in the test set.

Then, the design matrix for multivariate Bayesian classification in the training set is

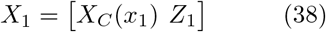

and the model assuming the *j*-th class for the *i*-th observation in the test set is

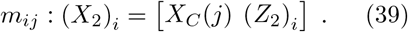

Similarly, the design matrix for multivariate Bayesian regression in the training set is

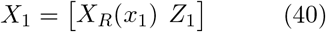

and the model assuming target value *λ* for the *i*-th observation in the test set is

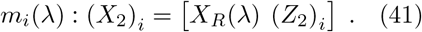

In other words, different than class indices and target values which are known in the training data and considered unknown in the test data, covariates will be considered known in both parts of the data set. Given these modified design matrices *X*_1_ and (*X*_2_)_*i*_, the process of calculating posterior probabilities is essentially the same, e.g. given by equation for MBC and (36) for MBR.

### 2.6 Cross-validation

Multivariate Bayesian inversion, as outlined in Sections 2.4.1 and 2.4.2, relies on cross-validation to predict labels based on features. Cross-validation (CV) successively splits a given data set into training and test sets and evaluates predictive performance on the test set, which is repeated until each data point has once been part of the test set [35, 36].

Unless otherwise stated, the following simulations (see Section 3) and analyses (see Section 4) use *k-fold cross-validation* with *k* = 10. This means that the data set is split into 10 equally large subsets (or sets as equally large as possible, if the number of CV folds *k* does not divide the number of data points *n*^1^) and in each CV fold, 9 subsets are used for training, i.e. estimation of the parameters (see Section 2.3.1), and the remaining subset is used for testing, i.e. identification of the labels (see Section 2.3.2). Splits into CV folds are non-random, but strictly sequential by index of data point.

For multivariate Bayesian classification (Simulations 1-3, Analyses 1-3), we use 10-fold CV *on points per class*. This means that all classes are split into 10 equally large subsets separately, such that in each CV fold, the distribution of classes is the same in training and test set (or as close as possible, if the number of CV folds does not divide the number of data points per class). This ensures that class imbalances are preserved in all subsets and thus do not have a spurious influence on classification.

For Analysis 2 (MNIST digit recognition; see Section 4.2) and Analysis 4 (brain age prediction; see Section 4.4), we did not use cross-validation, as the data sets were already partitioned in training and validation sets. In these cases, the posterior distribution over model parameters was estimated from the training set, the class labels (Analysis 2) or target values (Analysis 4) were predicted in the validation set and predictive performance measures are solely based on the validation set.

### 2.7 Comparison with other Bayesian methods

There exist a number of Bayesian approaches to machine learning [12] some of which are closely related to the general statistical approach presented here.

MBI differs from *Bayesian discriminative methods* such as Bayesian logistic regression [11, ch. 16] and Bayesian linear regression [11, ch. 14] by the fact that the latter typically model the predictive distribution of the labels given the features, i.e. the probability distribution *p*(*X*| *Y*) (cf. Section 2.1.1). MBI in turn is more closely related to *Bayesian generative methods* such as Gaussian naive Bayes classifiers [37] or Variational Gaussian mixture models [38] which describe the generative distribution of the features given the labels, i.e. the probability distribution *p*(*Y* |*X*), and later inverts this distribution to infer on labels given the features.

MBC and MBR are *multi-stage procedures*, meaning that data are splitted into training and test sets, in order to first estimate parameters of the generative model then extract values of the label variable (see Figures 1 and 2). This is in contrast to expectation maximization (EM) [39] and variational Bayesian (VB) [40] techniques which treat labels as latent random variables and jointly estimate the posterior distribution over label variables and model parameters.

The generative model underlying MBC is equivalent to the model behind (Bayesian) *linear discriminant analysis* (LDA) [41] when the design matrix only includes categorical regressors (*X* = *X*_*c*_(*x*)) and when all observations are regarded as independent (*V* = *I*_*n*_). In this case, all data points in the training set are assumed to follow multivariate normal distributions with class-specific means and class-independent covariance. By estimating *B* and *T* (cf. eq. 10), MBI implicitly estimates those class means and covariance matrix, such that new data points in the test set can be assigned to one class. This “LDA scenario” is exemplified in Simulation 3 (see Section 3.3) and Analyses 1/2 (see Sections 4.1/4.2).

The *novel contribution* here is that MBI generalizes (Bayesian) LDA (i) by using the same generative model for classification and regression, (ii) by allowing for correlations between data points in this model, (iii) by systematically integrating covariates into predictive analysis and (iv) by incorporating prior knowledge about class frequencies.

## 3 Simulations

In this section, we apply cross-validated multivariate Bayesian inversion to several synthetic data sets necessitating either classification (Simulations 1-3) or regression (Simulation 4). For each of these examples, we describe (i) the purpose behind the simulation study, (ii) the generative model underlying the data set, (iii) possible scientific interest in data sets of this form, (iv) steps of data analysis, we give a (v) summary of the results and provide (vi) a conclusion.

When of interest (Simulations 2-4), we compare MBI’s performance with that of support vector machines (SVM), as a representative easy-to-implement and currently popular alternative approach [42, 43].

### 3.1 Simulation 1: two-class classification

#### Purpose

The purpose of this simulation is to show how between-class variance (quantified as the Euclidean distance between class means) as well as within-class variance (quantified as the standard deviation around class means) influence classification accuracy in multivariate Bayesian classification (MBC).

#### Generative model

We simulate two classes of equal size in two dimensions: Each observation *y*_*i*_ ∈ ℝ^1×2^, *i* = 1, …, *n* is generated as

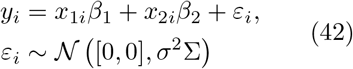

where *n* = 250 (with 125 per class) and some correlation between the two dimensions is induced via the between-feature covariance matrix

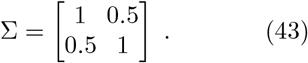

The terms *x*_1*i*_, *x*_2*i*_ ∈ *{*0, 1} are indicating the presence of the two classes and the class means are given by (see Figure 3)

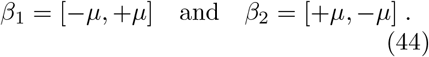

**Fig. 3.**
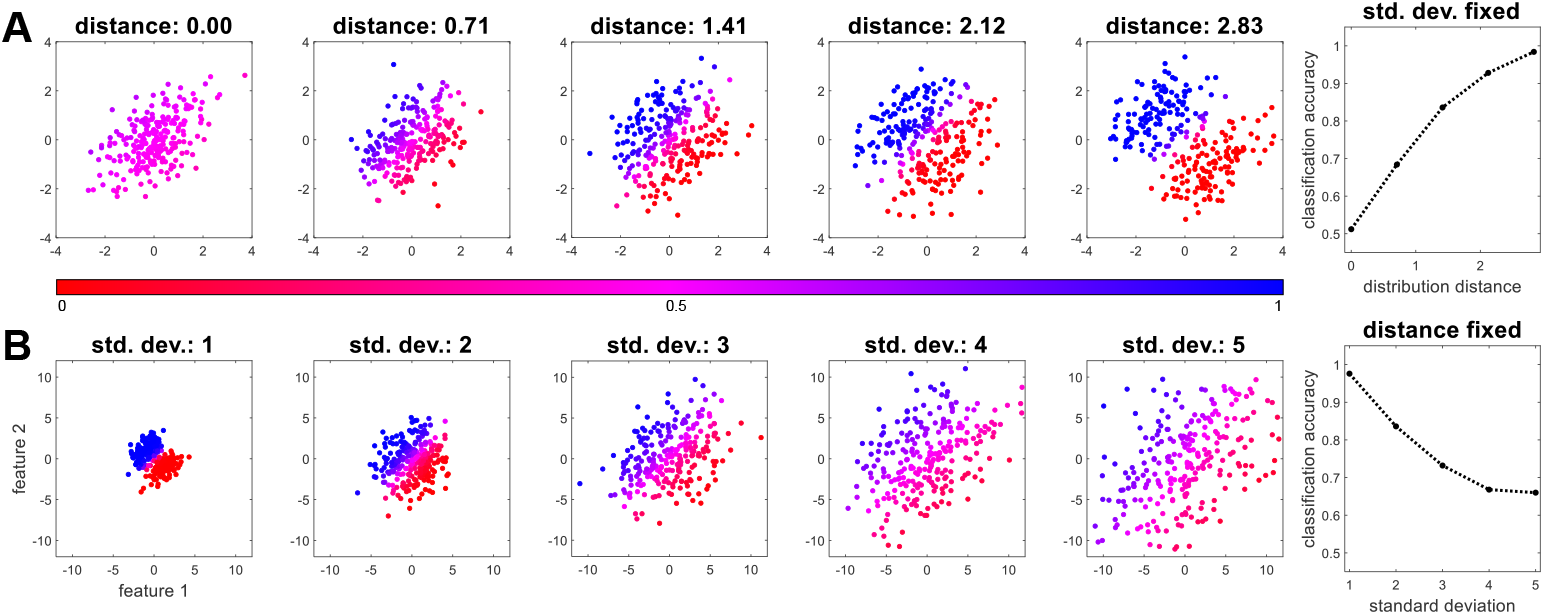
Simulation 1: two-class classification. Simulated data (*n* = 250) from two classes (blue, red) in two dimensions (x, y) when varying **(A)** the distance between the distributions (measured as Euclidean distance between the multivariate means) or **(B)** the spread of the distributions (measured as standard deviation of the noise). The color of each dot indicates the posterior probability assigned to this data point in multivariate Bayesian classification (see colorbar). Assigning classes by maximum posterior probability results in a classification accuracy for each of the scenarios (right panels).

Taken together, this is equivalent to the model behind linear discriminant analysis (LDA) [41], with class-dependent multivariate means and class-independent between-feature covariance. MBC generalizes LDA to additional variables influencing the features (see Simulation 2), an arbitrary number of classes (see Simulation 3) and potential between-observation correlation (see Simulation 4).

The parameters *µ* and *σ* are controlling (i) the distance between the centers of the two distributions and (ii) the amount of spreading (standard deviation) for the two distributions. In our simulations, they are specified as

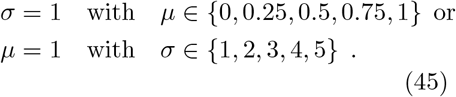

#### Scientific interest

Note that *µ/σ* is proportional to the Mahalanobis distance between the two distributions [44] which is a measure for class separability: With large *µ* and small *σ*, observations from the two classes are very distinct, whereas with small *µ* and large *σ*, the two classes are practically overlapping (see Figure 3A/B), which makes classification either a very easy or a very hard task.

Thus, a scientific question related to this simulated data set – or a real data set of the same form – could be how well the two classes can be separated, given some difference between class means and some variation within classes.

#### Data analysis

MBC is performed according to the methodology described in Sections 2.3.1, 2.3.2 and 2.4.1 based on the multivariate general linear model

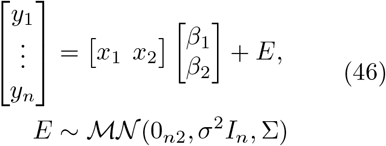

and using *k*-fold cross-validation (CV) with *k* = 10 CV folds.

#### Results

We observe that, trivially, (i) with increasing distance between the distributions, classification accuracy increases, and (ii) with increasing variance of the distributions, classification accuracy decreases (see Figure 3). Moreover, (iii) when the distance between distributions is zero, all points receive a posterior probability close to 1*/*2 (see Figure 3A), and (ii) when variance of distributions is large, many close to the decision boundary points receive such an indecisive posterior probability (see Figure 3B).

#### Conclusion

We have seen that classification accuracy in binary classification increases with distance of class means and decreases with variability of classes in multivariate space. Moreover, posterior class probabilities reflect the certainty of predictions made by MBC, with posterior probability becoming practically 1*/*2, if the two classes overlap.

### 3.2 Simulation 2: two classes with confound

#### Purpose

The purpose of this simulation is to show that including a confound (i.e. a covariate that systematically varies with class membership) into the classification model (MBC) results in higher classification accuracy than removing the covariate effect from the feature variables before classification (SVC).

#### Generative model

We again simulate two-dimensional data by drawing from two classes, but this time adding a continuous covariate to the outcome variables: More precisely, each observation *y*_*i*_ ∈ ℝ^1×2^, *i* = 1, …, *n* is generated as

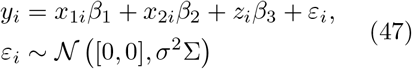

where *n* = 250, the class means *β*_1_ and *β*_2_ are as before (see Section 3.1) with *µ* = 1, *σ* = 2 and Σ given in (43). The continuous variable *z*_*i*_ ∈ [−1, +1] is sampled as

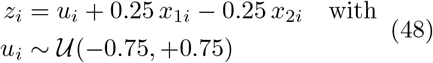

where 𝒰 (*a, b*) is a uniform distribution with minimum *a* and maximum *b*. This sampling causes *z*_*i*_ ∈ [−0.5, +1] for class 1 and *z*_*i*_ ∈ [1, +0.5] for class 2 (see Figure 4B), such that the covariate *z* is mildly correlated with class membership and tends to be higher for class 1. The effect of the covariate on the two features is specified as

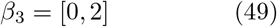

which means that the covariate has no effect on *y*_*i*1_ and exclusively acts on the second feature *y*_*i*2_ (see Figure 4B).

**Fig. 4.**
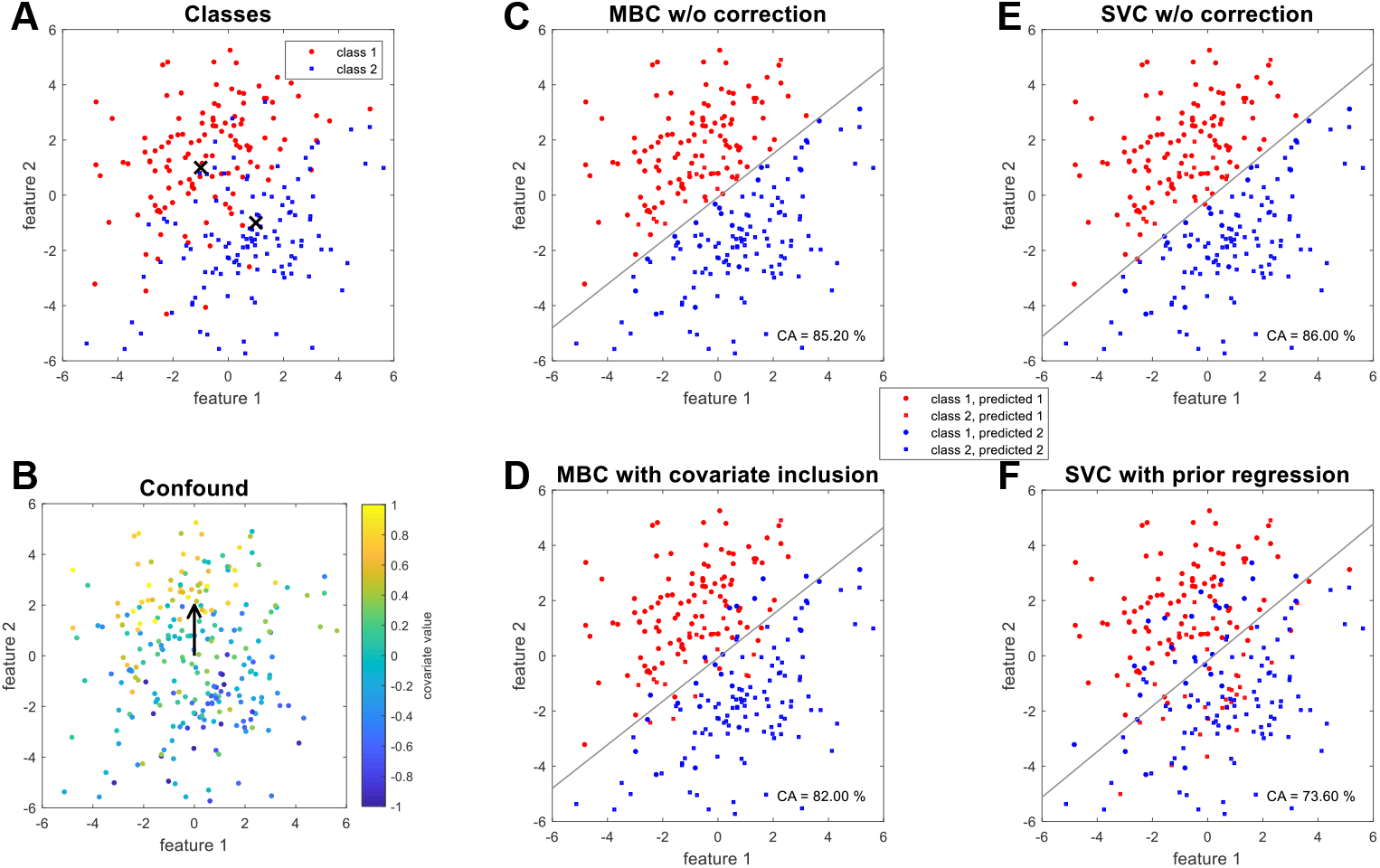
Simulation 2: two classes with confound. **(A)** Two-dimensional data (*n* = 250) is simulated as coming from two classes (class means indicated by black crosses) and **(B)** there is an additional effect of a confounding variable in one of the dimensions (covariate effect indicated by black arrow). **(C)** When performing MBC, classes can be separated with high accuracy (decision boundary shown as gray line). **(D)** Because this effect is in part based on the covariate, accounting for this confound reduces classification accuracy. **(E)** When performing SVC, a similar linear decision boundary also results in high accuracy. **(F)** Because accounting for the covariate is done via prior regression in SVC, the confounding nature of the covariate is not properly accounted for and classification accuracy decreases by much more. Also note that MBC as well as SVC with correction achieve classification beyond linear boundaries (see panels D and F), but for different reasons: MBC, because it includes an additional covariate into the model; SVC, because prior regression transforms the data into another space (for comparison purposes, data are plotted in the original space).

#### Scientific interest

When a data set contains a continuous variable that has an effect on the features, but also varies with class membership (such as *z*_*i*_), accounting for the effect of this confounder is crucial, such that patterns observed in the features may not be mistaken as being due to class, but as (partly) due to the covariate.

Thus, scientific questions related to this simulated data set – or a real data set of similar form – could be whether the two classes can be separated based on the features when accounting for the effect of the covariate associated with class membership, and whether separability differs depending on how the covariate is accounted for.

#### Data analysis

MBC is performed using the MGLM framework as before (see Section 3.1). In order to account for the correlated covariate (see Section 2.5), the multivariate linear model is extended to

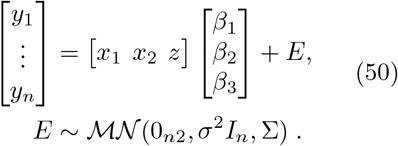

For comparison, SVC is performed by training a linear support vector machine with soft-margin parameter *C* = 1 on the data points from the two classes. In order to account for the confound in SVC, a linear regression model is fit to the features, removing the effect associated with *z* and keeping the residuals for classification. This is here referred to as the “prior regression” approach (see Figure 4F) and is known to be problematic sometimes [45, 46]. Both MBC and SVC are performed with 10-fold cross-validation.

#### Results

We observe that, the way the data are generated, values of the covariate are in fact correlated with the second feature and related to class membership (see Figure 4A/B). Both MBC and SVC result in linear decision boundaries separating the data points into predicted classes (see Figure 4C/E). Plausibly, because separation into classes is in part based on the covariate effect, accounting for this covariate results in a reduction of classification accuracy in MBC (see Figure 4D). However, because the SVM approach removes the covariate effect separately from classification and thus fails to account for the correlation of class membership and covariate, the classification accuracy reduces by much more in SVC (see Figure 4F).

In addition to this prior regression approach, we also tested a second correction method for SVC. This method includes the covariate as a feature, instead of removing its effect from the other features (see Section 4.4). We find that this preserves the classification accuracy of SVC without correction (CA = 86.00%; results not shown). However, this only shows that both, features and covariate, are correlated with class membership and it does not show how informative the features are about class membership when correcting for the effect of the covariate.

#### Conclusion

We have seen that covariate inclusion captures a confound’s effect on the features while accounting for the correlation of covariate and class which leads to a more realistic estimation of class separability than removing the confound’s effect via prior regression before classification.

### 3.3 Simulation 3: three-class classification

#### Purpose

The purpose of this simulation is to show that (i) MBC, unlike SVC, facilitates posterior class probabilities resulting in gradual classification boundaries and that (ii) prior class probabilities can be used to bias decisions into direction of particular classes by incorporating prior information.

#### Generative model

We simulate three classes of equal size in two dimensions: Each observation *y*_*i*_ ∈ ℝ^1×2^, *i* = 1, …, *n* is generated as

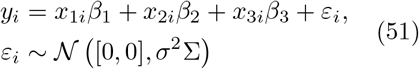

where *n* = 300 (with 100 per class), *σ* = 2, Σ is as given in (43) and *x*_1*i*_, *x*_2*i*_, *x*_3*i*_ ∈ {0, 1} where 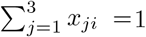 1. The class means are specified as (see Figure 5A)

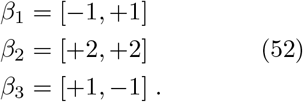

**Fig. 5.**
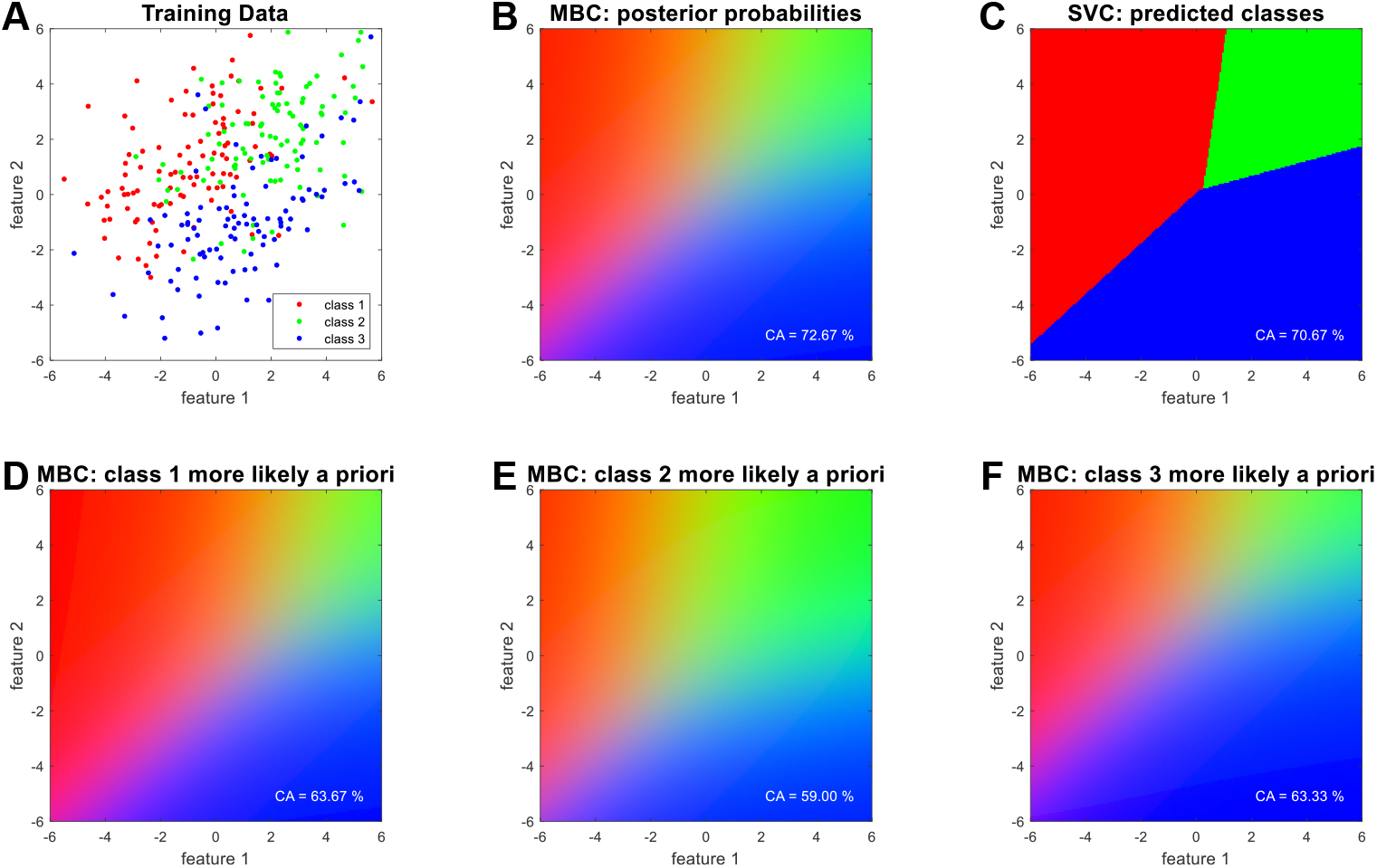
Simulation 3: three-class classification. **(A)** Simulated data (*n* = 300) from three classes (red, green, blue) in two dimensions (x, y). **(B)** Posterior class probabilities from MBC as a function of location in feature space are visualized as RGB triplets (e.g. [0.5, 0.0, 0.5] results in purple, which corresponds to equal posterior probability for classes 1 and 3; linear RGB converted to sRGB for visualization purposes). **(C)** Predicted classes from SVC are shown as a function of location in feature space. **(D)**,**(E)**,**(F)** Varying the prior class probabilities for MBC results in higher posterior probabilities for the class which is more likely a priori. Since classes were equally likely a priori, classification accuracy (CA) is highest when a uniform prior distribution is used (see panel B).

#### Scientific interest

Multi-class, or n-ary classification occurs frequently in empirical research problems such as optical character recognition, object classification or language detection. Thus, scientific questions related to this simulated data set – or a real data set of the same form – could be to what extent classes differ in multivariate space and whether some pairs of classes can be better distinguished than other pairs.

#### Data analysis

MBC is performed using the MGLM framework as before (see Section 3.1) based on the multivariate general linear model

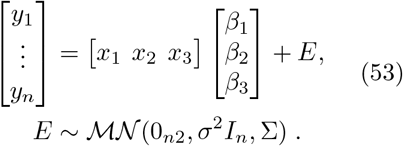

For exploratory reasons, we also perform MBC under manipulation of the prior class probabilities, i.e. by letting *p*(*m*_*ij*_) in (29) to be equal across classes, i.e. *p*(*m*_*ij*_) = 1*/*3 for all classes (see Figure 5B) or by setting *p*(*m*_*ij*_) = 2*/*3 for one class and *p*(*m*_*ij*_) = 1*/*6 for the other two classes (see Figure 5D/E/F).

For comparison, SVC is performed by training a linear support vector machine with soft-margin parameter *C* = 1 on the data points from the three classes. Both MBC and SVC are performed with 10-fold cross-validation.

#### Results

We observe that, when classification was performed by maximizing posterior probability, MBC and SVC would give rise to nearly identical decision boundaries (see Figure 5B/C). However, close to the decision boundaries, MBC provides graded information in the form of posterior probabilities rather than discrete predictions. Moreover, when incorporating prior knowledge by e.g. increasing the prior probability for one of the classes, the decision boundaries move accordingly, allocating more posterior probability to the respective class (see Figure 5D/E/F).

#### Conclusion

We have seen that MBC provides probabilistic information in multi-class decision problems and that it allows to incorporate prior information about class frequency, according to possible knowledge about the prevalence of categories.

### 3.4 Simulation 4: continuous regression

#### Purpose

The purpose of this simulation is to show that MBR, in the presence of between-observation covariance, achieves predictive correlation comparable with SVR and results in predicted targets whose distribution is more similar to the training data than with SVR.

#### Generative model

We simulate multi-dimensional data with continuous effects: Each observation *y*_*i*_ ∈ ℝ^1×*v*^, *i* = 1, …, *n* is generated as

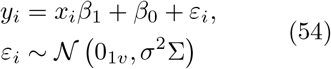

where *x*_*i*_ ∈ [−1, +1] is a continuous regressor sampled from a uniform distribution, *β*_1_ and *β*_0_ are 1 ×*v* row vectors with slopes and intercepts for the features *y*_*i*_ and Σ is a *v* ×*v* matrix inducing between-feature covariance^2^:

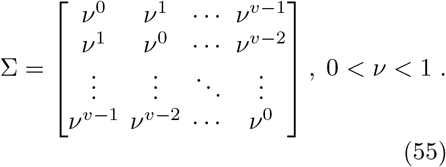

For demonstration purposes, we here also modulate the covariance between data points by applying an *n*× *n* matrix *V* inducing between-observation covariance:

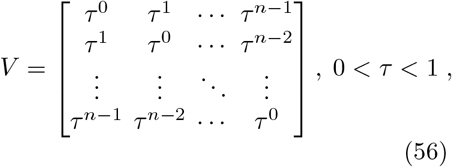

such that the simulation model for the entire data set is given by:

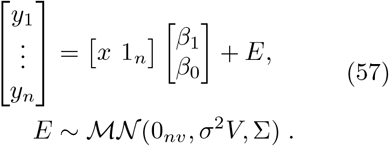

In our simulations, we set *n* = 200, *v* = 10, *σ* = 1, *ν* = 0.5, *τ* = 0.25 and all entries of *β*_0_ and *β*_1_ were sampled from the standard normal distribution 𝒩 (0, 1).

#### Scientific interest

If the rows of *Y* correspond to units of time (i.e. time series measurements), *τ* may also be called “time constant” governing the “temporal covariance” *V*. If the columns of *Y* correspond to units of space (e.g. measurement locations), then *ν* may also be called “space constant” governing the “spatial covariance” Σ [16]. In this situation, a scientific question could be whether and to what extent the continuous quantity *x*_*i*_ can be inferred based multivariate data *y*_*i*_, accounting for (known) temporal covariance and (estimated) spatial covariance. Moreover, values of *x*_*i*_ may fall into a range that may not necessarily preserved by a particular algorithm.

#### Data analysis

MBR is performed according to the methodology described in Sections 2.3.1, 2.3.2 and 2.4.2 based on the multivariate general linear model given by (57), with the aim of reconstructing the target values *x*_*i*_, *i* = 1, …, *n*. As the prior density *p*(*x*_*i*_) = *p*(*λ*) in (36), we use a continuous uniform distribution, i.e. *p*(*λ*) = 1*/*2 where −1 *< λ <* +1. Resulting posterior densities are calculated via discretization of the interval [−1, +1] in steps of 0.01 (see Figure 6D).

**Fig. 6.**
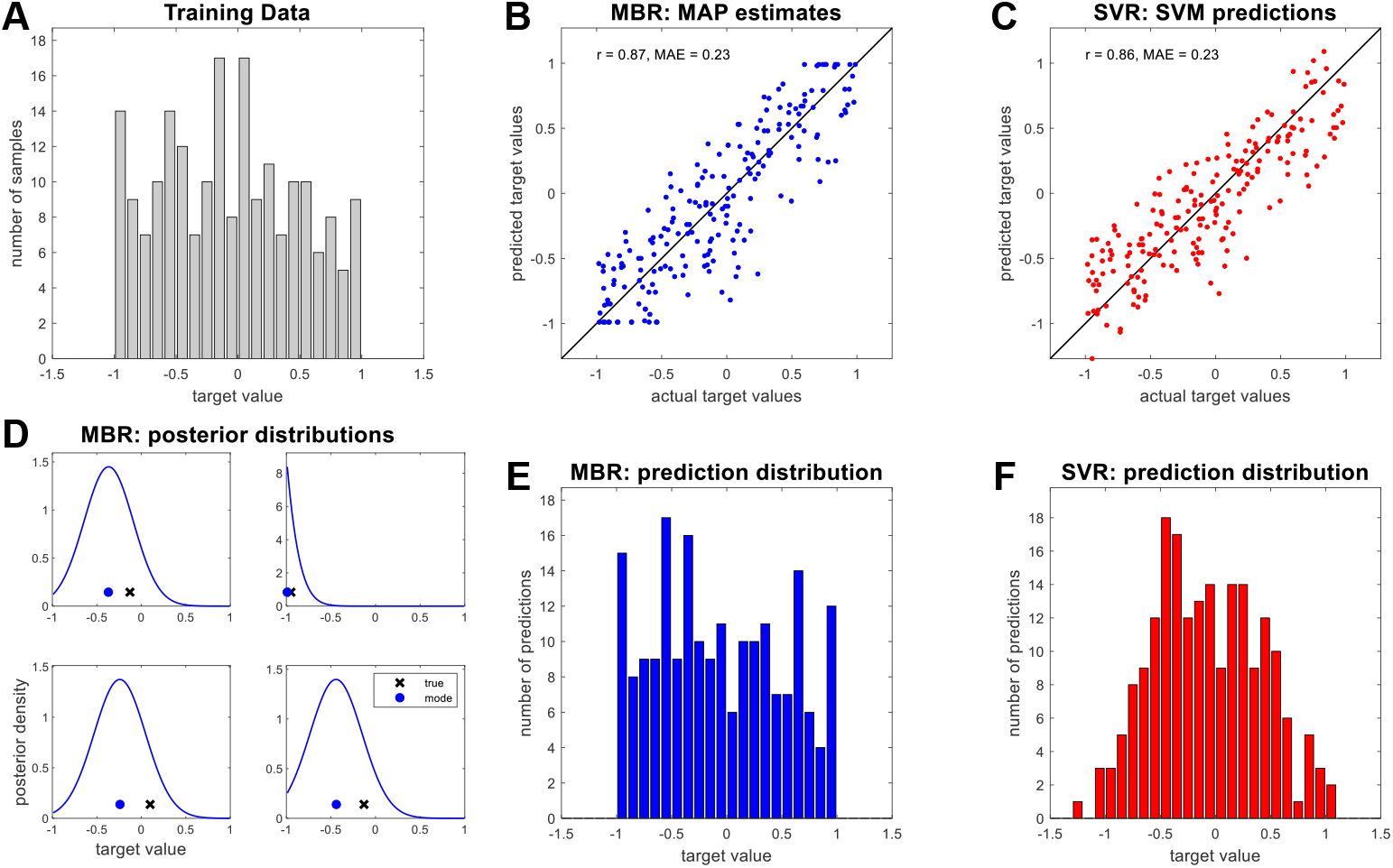
Simulation 4: continuous prediction. **(A)** A continuous predictor variable *x* (the histogram of which is shown) is used to sample a 200 × 10 data matrix *Y* (for details, see text). **(B)** Maximum-a-posteriori (MAP) estimates from MBR and **(C)** support vector machine (SVM) predictions from SVR are plotted against the actual labels. **(D)** Exemplary posterior target densities for the first four simulated data points, along with MAP estimate (blue dots) and true values (black crosses). **(E)** MAP estimates and **(F)** SVM predictions for the reconstructed targets are plotted as histograms.

For comparison, SVR is performed by training a linear support vector machine with soft-margin parameter *C* = 1 on the data matrix *Y* and the target vector *x*. Both, MBC and SVC are performed with 10-fold cross-validation.

#### Results

We observe that, trivially, maximum-a-posteriori (MAP) estimates from MBR fall into [−1, +1], because the prior probability is zero outside this interval. Notably, the resulting distribution of predicted target values is closer to the distribution of actual target values (see Figure 6A) for MAP estimates from MBR (see Figure 6E), when compared with SVM predictions (see Figure 6F). MBR and SVR are otherwise comparable in terms of predictive correlation and mean absolute error (see Figure 6B/C). In addition to this, it appears that posterior target densities *p*(*λ*| *y*_2*i*_) from MBR are univariate normal distributions when uniform prior densities are applied (see Figure 6D).

#### Conclusion

We have seen that MBR is able to reconstruct continuous labels from a multi-dimensional feature space and that it can effectively constrain the reconstructed variable into its natural range by a prior distribution restricted to that range.

### 3.5 Comparison with alternative approaches

#### Purpose

In addition to comparison with SVMs (see Figures 4-6), we also compare MBI’s simulation performance against that of other ML approaches for selected simulations. This includes generative and discriminative as well as Bayesian and non-Bayesian methods for classification and regression problems.

#### Data analysis

We re-analyze data from Simulation 3 using the following techniques:

- *naive Bayes classification* (GNB; multivariate normal likelihood, uniform prior);
- *linear discriminant analysis* (LDA; linear discriminant function);
- *multiclass logistic regression* (LogReg; regularization parameter *λ* = 1);
- *random forrest classification* (RFC; bootstrap aggregation)
- *neural network classification* (NNC; two hidden layers with ten neurons each, sigmoid activation function)

Moreover, we re-analyze data from Simulation 4 using the following techniques:

- *naive Bayes regression* (GNB; multivariate normal likelihood with linear effect of label variable on feature variables, uniform prior density over label variable);
- *multiple linear regression* (LinReg; weighted least squares estimation);
- *random forrest regression* (RFR; bootstrap aggregation)
- *neural network regression* (NNR; two hidden layers with ten neurons each, linear activation function)

We distinguish between generative approaches, i.e. methods modelling feature variables as a function of the label variable (e.g. LDA, and including MBC), and discriminative approaches, i.e. methods modelling the label variable as a function of the feature variables (e.g. LogReg, and including SVC).

In order to systematically compare diverse approaches, predictive performance of all techniques is measured as rank graduation accuracy (RGA) [47] which always falls between 0 and 1, with a value of 0.5 corresponding to chance-level performance. RGA is equivalent to area under the curve (AUC) in binary classification, but generalizes to n-ary classification and regression problems [48].

#### Results

Because MBC and LDA are equivalent for multi-class classification with i.i.d. observations and without covariates (see Section 2.7), both show identical performance for Simulation 3 (RGA = 0.8591). As expected, they perform better than GNB which disregards covariances between features (RGA = 0.8448). The highest performance is achieved by SVC (RGA = 0.8611), all other discriminative approaches have performance lower than that of MBC (see Table 3).

**Table 3.** Simulations: comparisons with alternative approaches. Predictive performance for generative and discriminantive techniques when analyzing data from Simulation 3 and Simulation 4. For details regarding MBC and SVC, see Figure 5, and for details regarding MBR and SVR, see Figure 6. In all cases, predictive performance is given as rank graduation accuracy (RGA) [48].

| Simulation 3 | RGA | Simulation 4 | RGA |
| --- | --- | --- | --- |
| *generative methods* |  |  |  |
| naive Bayes classification | 0.8448 | naive Bayes regression | 0.9218 |
| multivariate Bayesian classification | 0.8591 | multivariate Bayesian regression | 0.9327 |
| linear discriminant analysis | 0.8591 |  |  |
| *discriminative methods* |  |  |  |
| multiclass logistic regression | 0.8544 | multiple linear regression | 0.9250 |
| support vector classification | 0.8611 | support vector regression | 0.9309 |
| random forrest classification | 0.8169 | random forrest regression | 0.9143 |
| neural network classification | 0.7818 | neural network regression | 0.9318 |

In Simulation 4, differences between regression techniques are negligible, with MBR (RGA = 0.9327) and NNR (RGA = 0.9318) performing best among generative and discriminative approaches, respectively; GNB (RGA = 0.9218) and RFR (RGA = 0.9143) performing worst among generative and discriminative approaches, respectively. All other discriminative approaches have predictive performance between that of GNB and NNR (see Table 3).

#### Conclusion

MBI-based predictive analysis for classification and regression reaches predictive performance comparable with that of established (generative or discriminative, Bayesian or non-Bayesian) ML techniques in simulation studies.

## 4 Applications

In this section, we apply cross-validated multivariate Bayesian inversion to several empirical data sets necessitating either classification (Analyses 1-3) or regression (Analysis 4). Again, we describe (i) the purpose behind each empirical example, (ii) specificities, dimension (and possibly, acquisition) of the respective data set, (iii) possible scientific interest in this data set, (iv) steps of data analysis, we give a (v) summary of the results and provide (vi) a conclusion.

For all analyses, we compare MBI’s performance with that of support vector machines (SVM), as a popular ML approach that is often used for multivariate continuous data [42, 43].

### 4.1 Analysis 1: Egyptian skull data

#### Purpose

The purpose of this analysis is to show that MBC achieves (above-chance) classification accuracy comparable with SVC in different binary and multi-class classification scenarios for a single data set without covariates.

#### Data set

The Egyptian skull data set ([49]; [26, p. 287]) comprises 4 skull measurements (maximal breadth of the skull, MB; basibregmatic height, BH; basialveolar length, BL; nasal height, NH) from 5 × 30 human skulls coming from 5 time periods (4000 BCE, 3300 BCE, 1850 BCE, 200 BCE, 150 CE). Thus, we have *v* = 4 features from *n* = 150 observations belonging to *C* = 5 classes. The entire data set is visualized in Figure 7A.

**Fig. 7.**
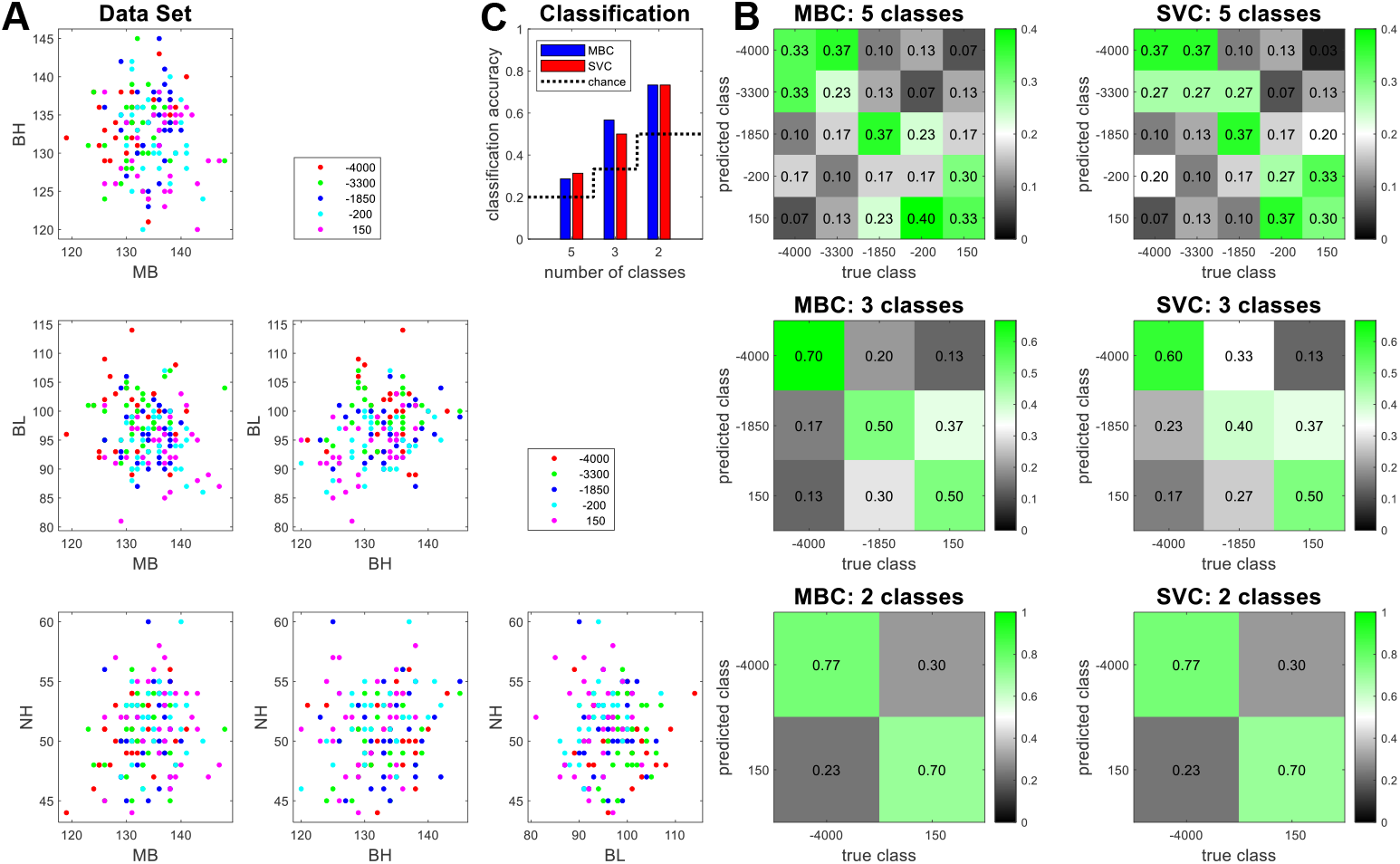
Analysis 1: Egyptian skull data. **(A)** Maximal breadth of the skull (MB), basibregmatic height (BH), basialveolar length (BL) and nasal height (NH) are shown for 30 skulls each from around 4000 BCE (red), 3300 BCE (green), 1850 BCE (blue), 200 BCE (cyan) and 150 CE (magenta). Note that all measurements are given in integer millimeters (mm), causing an impression of discreteness in the plots. **(B)** 5-class, 3-class and binary classification were exercised with the goal to detect the historical era based on the four skull measurements. Confusion matrices show the proportion of predicted classes (y-axis), given the true class (x-axis). **(C)** Classification accuracy, i.e. the proportion of correct predictions across all observations, is shown as a function of classification algorithm and number of classes. Classification accuracy decreases with increasing number of classes, but MBC and SVC achieve comparable performance.

#### Scientific interest

Scientific questions related to this data set could be:

- To what extent are skulls from various eras differing in terms of MB, BH, BL, NH?
- Are skulls differing more strongly, if they are from temporally more distant eras?
- Given a random skull, with what accuracy can it be assigned to a time period?

#### Data analysis

Analyses are performed for classification into five groups (all time periods) or distinction into three categories (−4000 vs. −1850 vs. 150) or binary classification (− 4000 vs. 150). MBC is performed as described in Section 2.4.1, setting *V* = *I*_*n*_ for no covariance between observations. SVC is performed, setting the soft-margin parameter *C* = 1. Both MBC and SVC are performed with 10-fold cross-validation.

#### Results

We observe that, as could be expected, classification accuracy increases with decreasing number of classes (see Figure 7C). However, MBC as well as SVC both exceed chance level in each classification task (see Figure 7C). Confusion matrices suggest that the most extreme time periods (−4000, 150) were predicted with highest accuracy (see Figure 7B) and that the 3rd era (−1850) can be better recognized than the 2nd and the 4th era (−3300, −200), possibly because the latter were closer in time to the 1st and the 5th era (−4000, 150) than to the middle-most time period (see Figure 7B).

#### Conclusion

We have seen that a comparably large number of classes (5) can be separated based on a comparably small number of samples per class (30) using a comparably low number of features (4). Moreover, confusion matrices show that misclassifications become less frequent with increasing temporal distance between true and predicted class which highlights empirical plausibility of the results.

### 4.2 Analysis 2: MNIST digit recognition

#### Purpose

The purpose of this analysis is to show that MBC (and SVC) accuracy is increasing with growing number of training samples and that MBC-based posterior probabilities approximate true class frequencies of predicted test data points.

#### Data set

The MNIST database [50, 51] consists of images of handwritten digits 0 to 9. Original images from the US National Institute of Standards and Technology (NIST) were size-normalized into a 20 ×20 pixel box, converting binary black-and-white images into gray-scale levels in the interval [0, 255], and placed into a 28 ×28 pixel image by computing the center of mass of the pixels. The data set contains *n*_1_ = 60,000 training samples, *n*_2_ = 10,000 test samples, the number of classes is *C* = |{0, …, 9}| = 10 and the number of features is *v* = 28 × 28 = 784.

#### Scientific interest

Scientific questions related to this data set could be:

- With what accuracy can handwritten digits be recognized from pixelated images?
- Which pairs of digits are more often confused with each other than other pairs?
- Given a digit image *Y* has been assigned a probability *p* of being digit *X*, what is the frequency of this assignment being correct?

#### Data analysis

During preprocessing, 4 pixels were removed on either side of each image, extracting only the center 20 ×20 pixels, resulting in a reduced number of features *v* = 20 ×20 = 400. Moreover, all pixel intensity values were rescaled into the interval [0, 1]. Each preprossed image matrix 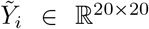 was vectorized into a row vector *y*_*i*_ ∈ ℝ^1×20·20^, giving an *n*_1_ × *v* training data matrix *Y*_1_ and an *n*_2_ × *v* test data matrix *Y*_2_.

MBC is performed as described in Section 2.4.1, setting *V* = *I*_*n*_. SVC is performed, setting the soft-margin parameter *C* = 1. Both MBC and SVC are performed for classification into ten categories (digits 0-9) and with 10-fold cross-validation.

#### Results

We observe that classification accuracy on the test set depends on the number of training samples used for estimating model parameters and that SVC reaches higher classification accuracy than MBC across all numbers of training samples, with both clearly exceeding chance level in the digit classification task (see Figure 8A). This is also evident from confusion matrices which show higher misclassification rates for MBC than for SVC, as the former especially confuses 3 with 5 and 4 with 9 (see Figure 8B/C).

**Fig. 8.**
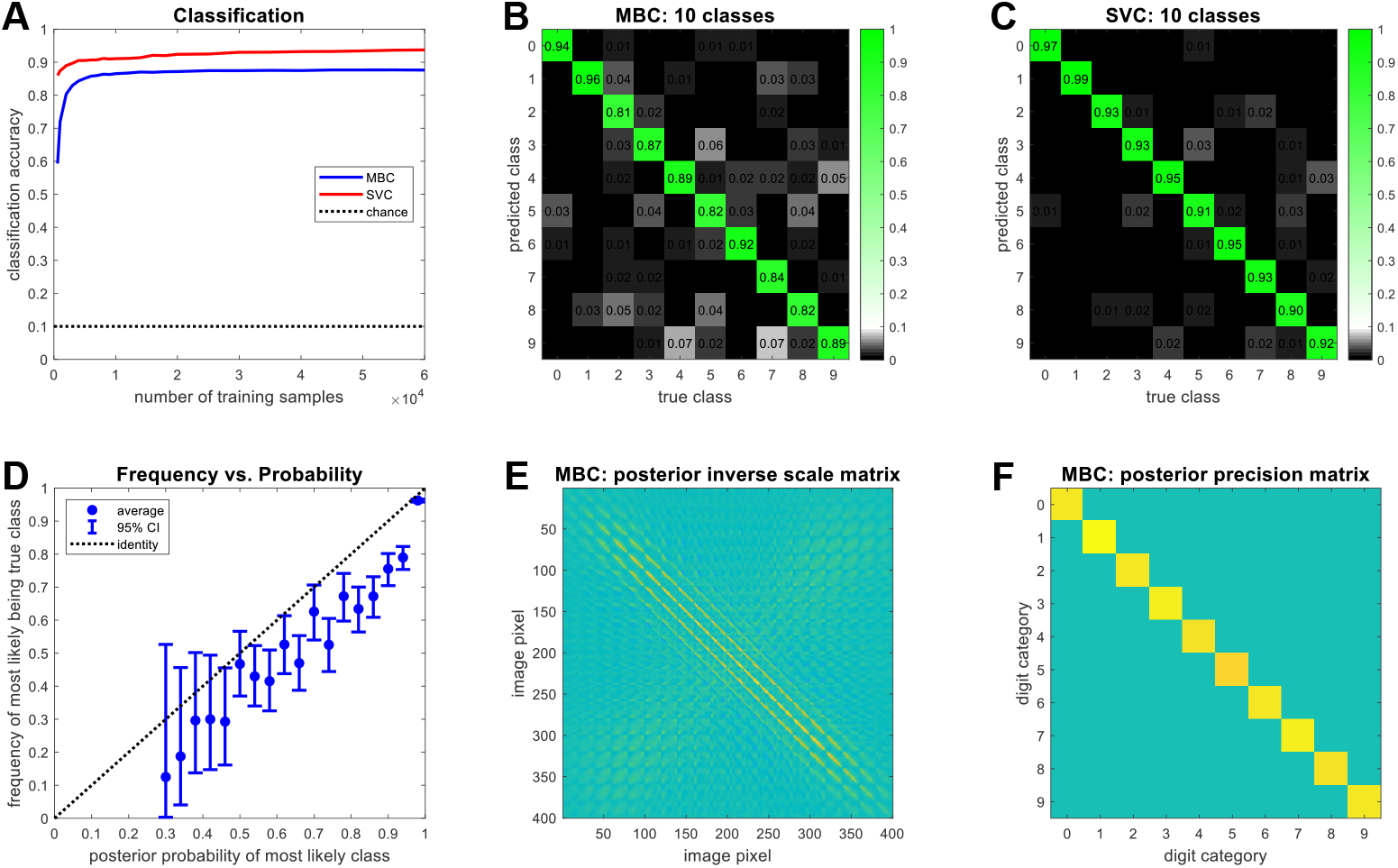
Analysis 2: MNIST digit recognition. **(A)** Classification accuracy in the test set (*n*_2_ = 10,000) is shown as a function of the number of training samples (ranging from 600 to 60k) used to estimate parameters for multivariate Bayesian classification (MBC; blue) and support vector classification (SVC; red). Classification analyses were performed to distinguish digits 0-9. For the full training set (*n*_1_ = 60,000), confusion matrices show the proportion of predicted classes (y-axis) given the true class (x-axis) for **(B)** MBC and **(C)** SVC. Misclassification rates are higher for MBC. **(D)** MBC assigns posterior class probabilities to test set samples and the optimal choice is given by predicting the class with the highest posterior probability. We plot the frequency of the most likely class being the actual class against binned highest posterior probability. Blue dots represent estimated frequencies and error bars denote 95% binomial confidence intervals. **(E)** MBC’s posterior inverse scale matrix Ω_1_ (after training), indicating covariance between image pixels. Nearby pixels shower higher covariance than pixels which are far apart. **(F)** MBC’s posterior precision matrix Λ_1_ (after training), indicating covariance between classes. Because classes are mutually exclusive, no between-class covariance exists.

**Fig. 9.**
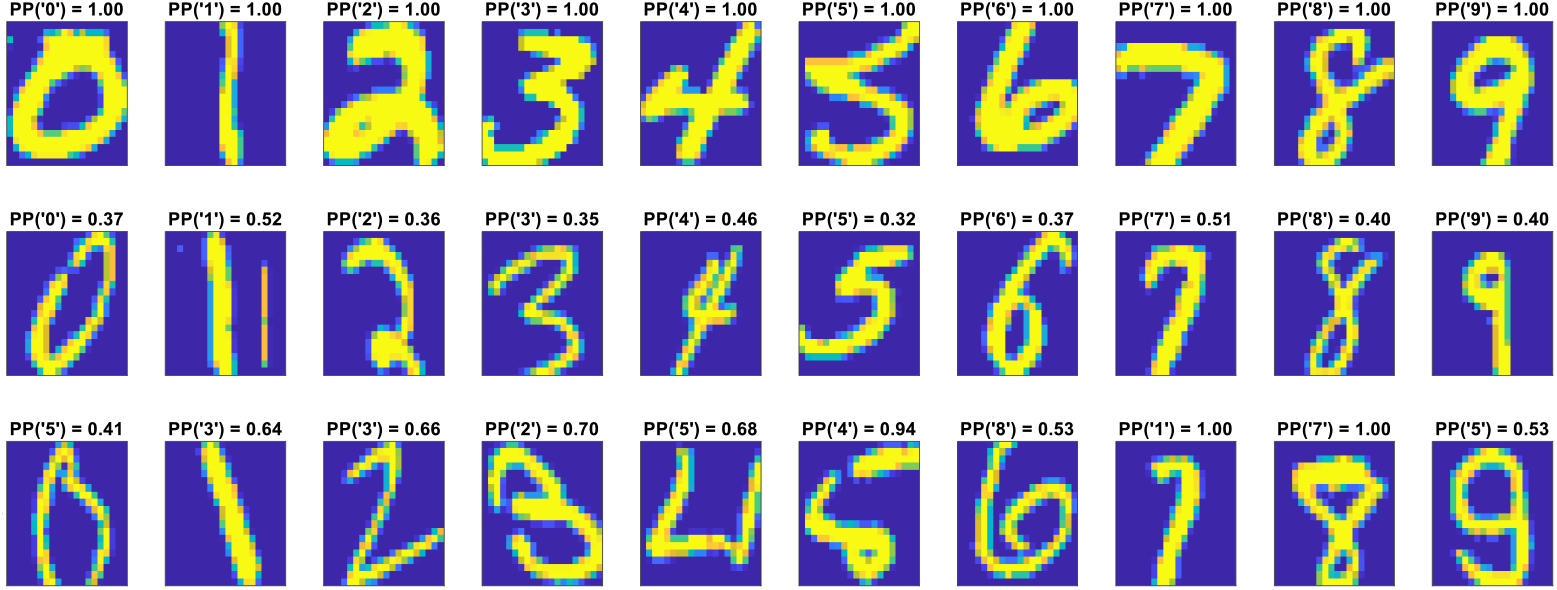
Analysis 2: exemplary test set cases. For each class of digits, three samples from the test set are shown with the highest posterior class probability assigned to these samples (“PP(‘5’) = 0.41” means that this test sample is most likely digit 5, with a posterior probability of around 41%). The first row shows correct classifications with the highest posterior class probability for each digit (“high-confidence hits”). The second row shows correct classifications with the lowest posterior class probability for each digit (“low-confidence hits”). The third row shows one random incorrect classification in which the class with the highest posterior probability differs from the true class (“misses”).

Next, we evaluated how accurately the posterior class probabilities calculated for test samples reflect the actual class of those samples. Each test sample is assigned ten posterior class probabilities (for digits 0-9) and can then be predicted as belonging to the class for which posterior class probability is highest. Therefore, we binned test data items by highest posterior probability (25 bins, bin width 0.04) and calculated, for each bin, the frequency that the class with the highest posterior probability was also the true class. There is a linear-increasing relationship between posterior probability of the most likely class and the frequency of the most likely class being the true class (see Figure 8D), indicating that posterior probabilities are somewhat veridical. However, the relationship is slightly below the identity line (see Figure 8D), suggesting that MBC posterior probabilities based on the MNIST training data are a bit overconfident.

For illustration, we also show the posterior inverse scale matrix Ω_1_ (cf. eq. 19c), embodying the covariance between features (see Figure 8E), and the posterior precision matrix Λ_1_ (cf. eq. 19b), embodying the covariance between classes (see Figure 8F), according to the training data. As nearby image pixels tend to be correlated (digits are contiguous, so neighboring pixels have similar intensity values) and all image matrices were vectorized before training, Ω_1_ is characterized by diagonal lines which become weaker, the higher the distance from the main diagonal (and thus, the more two pixels are apart from each other in the image). As classes are mutually exclusive (no data point belongs to two or more digits at the same time), Λ_1_ is a diagonal matrix with all off-diagonal entries (representing between-class covariances) being equal to zero.

Finally, we present exemplary test set cases which were correctly classified with either highest posterior probability (see Figure 12, top row) or lowest posterior probability (see Figure 12, middle row) or which were misclassified, with posterior probability being maximal for the wrong digit (see Figure 12, bottom row). These examples anecdotally show that images are more likely to be correctly classified, if they are drawn in bold, rather than thin lines (see Figure 12, top and middle row), and that they are more likely to be misclassified, if they are similar to other digits (see Figure 12, bottom row).

#### Conclusion

We have seen that classification accuracy improves with increasing size of the training set, as the posterior distribution over MGLM model parameters (see Section 2.2.1) is more accurately estimated when more training data are available. Moreover, posterior probabilities derived from MBC are veridical with respect to the true class in the sense that the they approximately reflect the frequency with which a test set sample is the class to which the highest posterior probability was assigned.

### 4.3 Analysis 3: birth weight data

#### Purpose

The purpose of this analysis is to show that MBC achieves higher classification accuracy than SVC and less “classification artefacts” (i.e. (near-)exclusive prediction of one class) in situations where one categorical variable (here: smoking) is predicted while other categorical variables (here: ethnicity) and parametric variables (here: age) influence the feature variables.

#### Data set

The low birth weight data set ([52]; [26, p. 288]) consists of a number of measurements from *n* = 189 United States mother-baby pairs which was collected to investigate potential causes of low birth weight. Collected variables include the birth weight (in kg), the mother’s weight (in kg), smoking status (non-smoker, smoker), ethnicity (white, black, other), hypertension (no, yes), the mother’s age (in yrs) and the number of visits to the doctor before birth. The entire data set is visualized in Figure 10A.

**Fig. 10.**
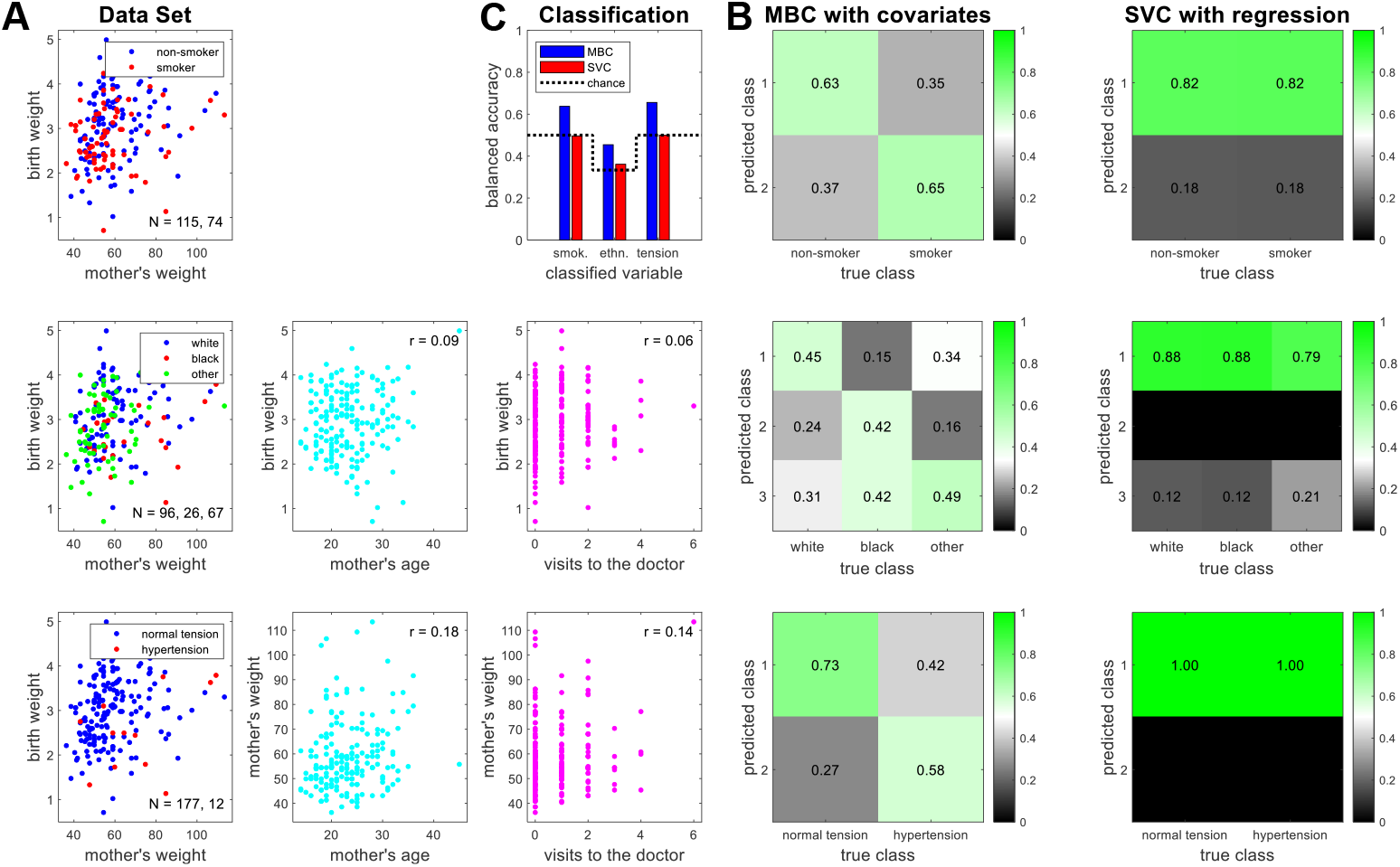
Analysis 3: birth weight data. **(A)** Birth weight [kg] and mother’s weight [kg] as a function of smoking status (top), ethnicity (middle), hypertension (bottom) and as a function of mother’s age [yrs] (cyan) and visits to the doctor (magenta). **(B)** Classification analyses were executed with the goal to distinguish the discrete variables based on the two features, accounting for covariates. Confusion matrices show the proportion of predicted classes (y-axis), given the true class (x-axis). **(C)** Balanced accuracy, i.e. the average over class-wise accuracies (diagonal entries in confusion matrices), is shown as a function of classification algorithm and classified variable. Whereas SVC remains at chance level, MBC exhibits above-chance prediction accuracy.

#### Scientific interest

Presumably, the original purpose of acquiring the data set was to identify causes of low birth weight, i.e. to predict birth weight from the other variables. Turning this question around, one can also ask whether the the effects of lifestyle, demographics and health on body weight are so strong that these categorical variables can be detected from the parametric ones. In our analysis, the goal therefore is to predict the 3 discrete variables (smoking, ethnicity, hypertension) from the 2 continuous features (birth weight, mother’s weight). Challenges in this prediction task are (i) strongly unequal class sizes (see Figure 10A), (ii) controlling for the dimensions not currently predicted and (iii) confounding variables (mother’s age, visits to the doctor).

#### Data analysis

Analyses are performed for classification into two categories (smoking, hypertension) or three categories (ethnicity). MBC is performed as described in Section 2.4.1, setting *V* = *I*_*n*_ as data points were regarded independent. SVC is performed, setting the soft-margin parameter *C* = 1. Both MBC and SVC are performed with 10-fold cross-validation, as described in Section 2.6.

In order to account for potentially confounding variables, a design matrix *Z* was generated by forming difference regressors (+1 vs. − 1) for the categories not currently classified (e.g. hypertension when predicting smoking) and including continuous covariates (e.g. mother’s age) as further regressors. For MBC, this covariate design matrix *Z* was added to the categorical design matrix *X*_*C*_ used for classification itself, as described in Section 2.5. For SVC, a linear regression model is fit to the features, removing the effect associated with *Z* and keeping the residuals for classification analyses.

To assess classification performance, we used the balanced accuracy (BA), i.e. the average of class-wise accuracies, rather than the classification accuracy (CA), as CA can lead to misleading conclusions in the case of unequal group sizes which are however compensated by the BA [53, 54].

#### Results

We observe that SVC with prior regression fails to accomodate the unequal group sizes in all three classification tasks (smoking, ethnicity, hypertension), frequently predicting the largest class, independent of the true class (see Figure 10B). Note that, when samples were weighted by class size in the respective training set, predicted class was more correlated with true class for all three label variables, producing above-chance accuracies (results not shown). We here report unweighted results for SVM analyses, as no classification approach was informed with group sizes in training or test set analysis.

In contrast to SVC, MBC with covariate inclusion can seemingly adjust for the unequal numbers of samples, resulting in acceptable class-wise accuracies throughout (see Figure 10B). As a consequence, balanced accuracy based on SVC is at chance level whereas MBC achieves above-chance prediction accuracy (see Figure 10C). We hypothesize that MBC with covariate inclusion manages to account for interactions between discrete variables and for the effects of covariates (see Figure 10A), thereby more faithfully estimating the true pattern underlying the categories currently classified.

We also ran another SVC analysis that includes the covariates as features, instead of removing their effects from the other features (see Section 4.4). We find that this gives above-chance MBC-like balanced accuracies for smoking (BA = 65.29%) and ethnicity (BA = 48.12%), but chance-level performance for hypertension (BA = 50.00%; results not shown). This however only demonstrates that features and covariates *together* are predictive of smoking and ethnicity, but it makes no statement about how informative the features are about those variables when *correcting* for the covariates’ effects.

#### Conclusion

We have seen that MBC outperforms SVC in a relatively small data set with few features, potentially confounding continuous variables (here: mother’s age, visits to the doctor), potentially interacting discrete variables (here: smoking, ethnicity, hypertension) and highly unequal class sizes.

### 4.4 Analysis 4: brain age prediction

#### Purpose

The purpose of this analysis is to show that MBR achieves higher predictive correlation than SVR when variables covarying with the features are included as covariates influencing the features (MBR), rather than as features predicting the labels (SVR).

#### Data set

In brain age prediction, the goal is to estimate an individual person’s chronological age (in years) from structural magnetic resonance imaging (MRI) scans of this person [55, 56]. For the present analysis, we are working with dimensionality-reduced structural MRI information from *n* = 3,300 subjects that was obtained by averaging within the regions from an anatomical atlas of the human brain and already used in an earlier study [57].^3^

As the data were analyzed within a predictive analytics competition^4^, the analysis was not fully cross-validated, but the the data set was split into *n*_1_ = 2,640 training samples (available during the competition) and *n*_2_ = 660 validation samples (target values not disclosed, used to decide the competition) with chronological age between 17 and 89 years (see Figure 11, left column). For each of these subjects, gray matter (GM) and white matter (WM) densities for 116 brain regions were extracted. Additionally, each subject’s gender (coded as +*/−*1) and the site at which they were acquired (1 out of 17) were known [57, 58].

**Fig. 11.**
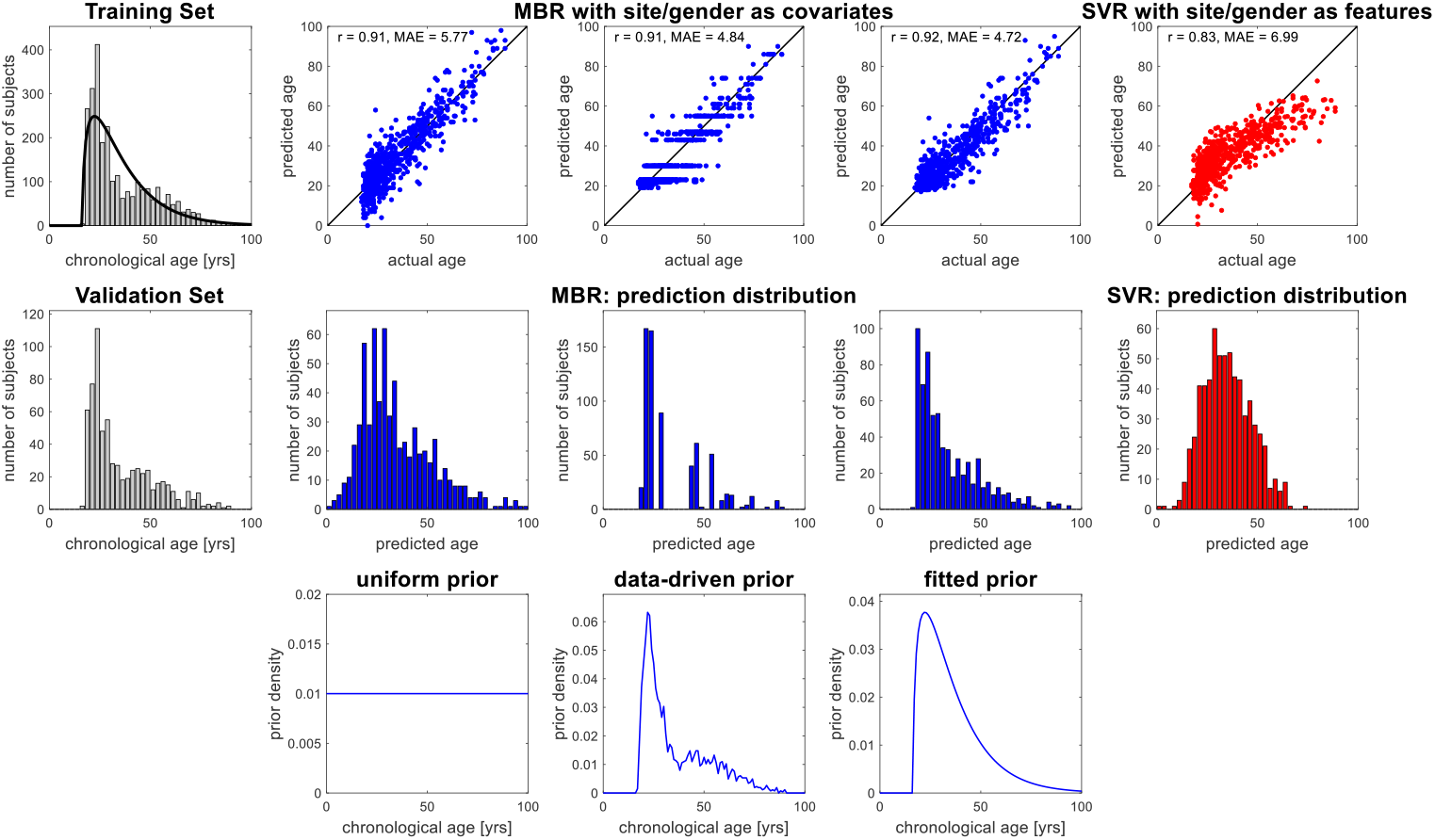
Analysis 4: brain age prediction. Chronological age in training and validation set (left, gray) was predicted from structural MRI features (not shown) using MBR with site and gender as covariates (middle, blue), using different prior densities on age (bottom row), and using SVR with site and gender as features (right, red), resulting in higher predictive correlation and lower absolute errors for MBR (top row) and distributions of predicted values that are closer to the actual distribution (middle row).

**Fig. 12.**
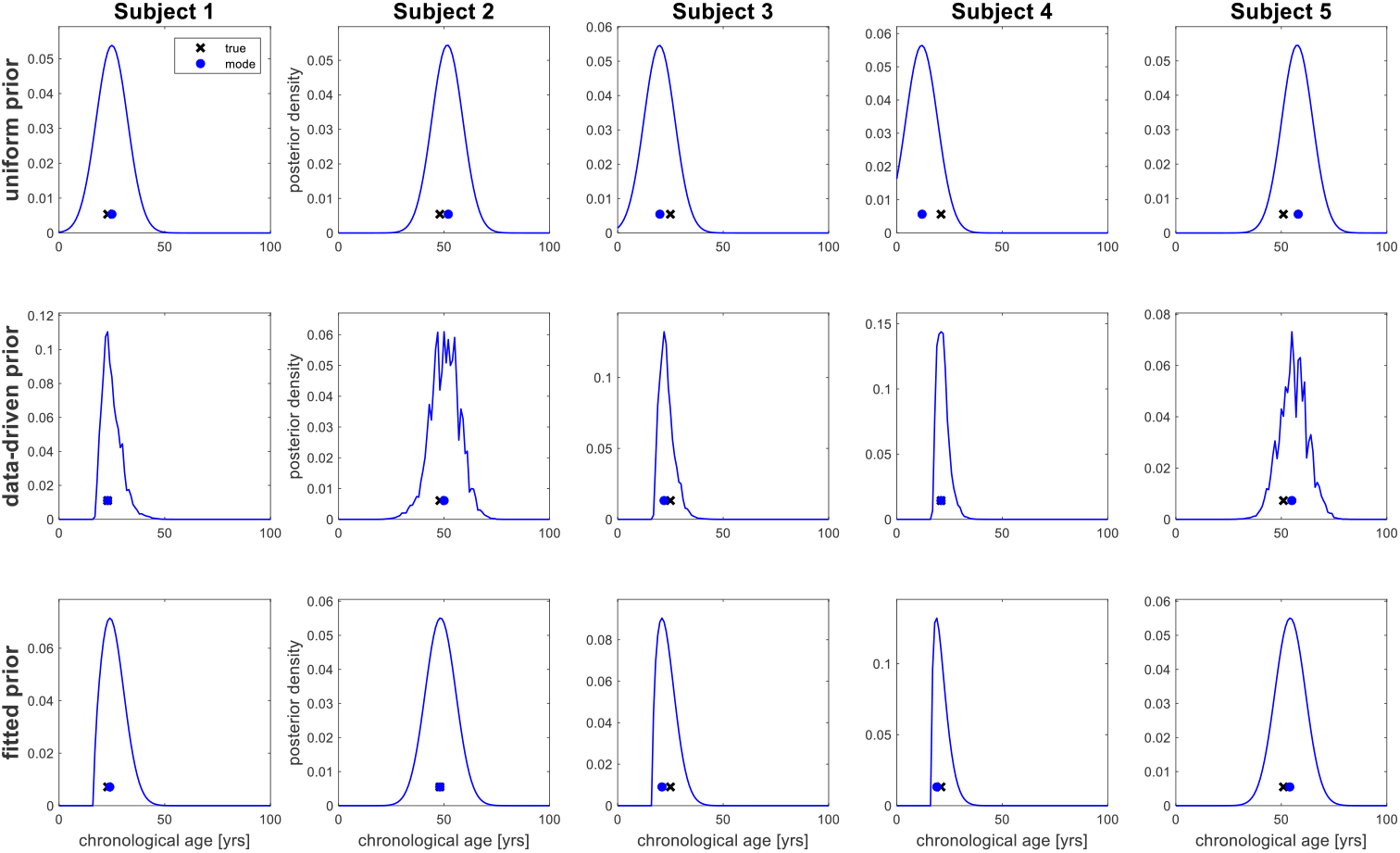
Analysis 4: posterior distributions. Exemplary posterior distributions over chronological age with maximum-a-posteriori estimates (blue dots) and true values (black crosses) for the first five subjects in the validation set (see Figure 11, left column) when using multivariate Bayesian regression with uniform prior (top row), data-driven prior (middle row) or fitted prior (bottom row) over chronological age (see Figure 11, bottom row). For explanations, see text.

#### Scientific interest

Scientific questions related to this data set could be:

- Which are the brain features that are most strongly varying with chronological age?
- With what precision can chronological age be detected based on brain features?
- How can the non-uniform distribution of target values in the training set be handled?

#### Data analysis

SVR was performed by calibrating a linear support vector machine with soft-margin parameter *C* = 1 on the training data, using chronological age as the target variable *x* and the 2 ×116 + 17 + 1 = 250 variables (= GM/WM × regions + sites + gender) as feature matrix *Y*, and then predicting chronological age in the validation data (see [57, sec. 2.2, 2.3.2] for details). Adding site and gender to the feature matrix, although different from the “prior regression” approach used above (see Simulation 2 and Analysis 3), is compatible with common practice in biomedical research in which covariates are added to the feature matrix when predicting class labels [59] or target values [60].

By contrast, in a statistical modelling context, it is not really useful to employ covariates such as site and gender as features for prediction of age, since gender was putatively controlled when splitting the data and the whole age range was putatively sampled at each site [61]. And even if this wasn’t the case, prediction of a subject’s chronological age from gender or site is not really something that would be demanded. Instead, structural MRI features could be varying as a function of gender or site, such that these must be included as influential covariates, not additional features.

MBR was therefore performed using the MGLM framework outlined above (see Sections 2.4.2 and 2.5) based on the multivariate general linear model

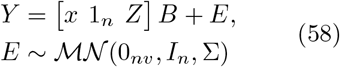

where *Y* is a *n*_1_ × (2 × 116) matrix of WM and GM densities, *x* is an *n*_1_ × 1 vector of chronological age, 1_*n*_ is a constant regressor of the same size and *Z* is a *n*_1_ ×17 covariate design matrix including indicator regressors for site and gender effects, such that *B* is a (2+17) × (2 ×116) matrix of regression coefficients, describing the influences of *x* and *Z* on *Y*. In this way, effects of site and gender on structural MRI measures are accounted for when reconstructing chronological age from these quantities.

For MBR, we applied three different prior target densities (see Figure 11, bottom row): (i) a uniform prior, with constant prior density for ages between 0 and 100 years; (ii) a data-driven prior, the probability for each integer age being equal to its frequency of occurence in the training set; and (iii) a fitted prior, from fitting a shifted gamma distribution to the target values in the training set (see Figure 11, top left). Like in Simulation 4, posterior densities we obtained via discretization of the interval [0, 100] in steps of 1.

#### Results

We observe that (i) all MBR variants achieve a higher predictive correlation (uniform: *r* = 0.91; data-driven: *r* = 0.91; fitted: *r* = 0.92) than SVR (*r* = 0.83) when predicting chronological age (see Figure 11, top row); (ii) this is underscored by the fact that the distribution of MBR predictions is closer to the actual distribution of targets than the distribution of SVR predictions (see Figure 11, middle row); and (iii) using the fitted prior density for age results in the smallest mean absolute error (MAE = 4.72 yrs), outperforming the uniform prior (MAE = 5.77 yrs) by about a year and SVR (MAE = 6.99 yrs) by more than two years (see Figure 11, top row).

We also present exemplary posterior target densities using the three different priors (see Figure 12) which anecdotally show how the data-driven and the fitted prior can occasionally draw the MAP estimates closer to the true value (see Figure 12, right column). Replicating Simulation 4, we also repeat our observation that posterior target densities from MBR seem to approximate univariate normal distributions when uniform prior densities are applied (see Figure 12, first row).

#### Conclusion

We have seen that MBR outperforms SVR by including confounding variables as covariates influencing the features, rather than as features allowing to predict the targets. Additionally, different prior distributions can be incorporated, with an ecologically plausible prior distribution resulting in the smallest mean absolute error.

### 4.5 Comparison with alternative approaches

#### Purpose

In addition to comparison with SVMs (see Figures 7-8, 10-11), we also compare MBI’s empirical performance against that of other ML approaches for selected analyses. This includes generative and discriminative as well as Bayesian and non-Bayesian methods for classification and regression problems.

#### Data analysis

We analyze data from Analysis 2 using the same techniques and settings described in Section 3.5: *naive Bayes classification* (GNB); *linear discriminant analysis* (LDA); *multiclass logistic regression* (LogReg); *random forrest classification* (RFC); *neural network classification* (NNC; one hidden layer with 400 neurons). Moreover, we re-analyze data from Analysis 4 using the same techniques and settings described in Section 3.5: *naive Bayes regression* (GNB); *multiple linear regression* (LinReg); *random forrest regression* (RFR); *neural network regression* (NNR; one hidden layers with 250 neurons).

As before, we distinguish between generative approaches (e.g. GNB, and including MBC) and discriminative approaches (e.g. RFC, and including SVC) and measure predictive performance using the rank graduation accuracy (RGA) [47] which always falls between 0 and 1, with a value of 0.5 corresponding to chance-level performance.

#### Results

Because MBC and LDA are equivalent for multi-class classification with i.i.d. observations and without covariates (see Section 2.7), both show identical performance for Analysis 2 (RGA = 0.9396). As expected, they perform better than GNB which disregards covariances between features (RGA = 0.8934). The highest performance is achieved by NNC (RGA = 0.9908) and all discriminative approaches except for LogReg have performance higher than that of MBC (see Table 4).

**Table 4.**
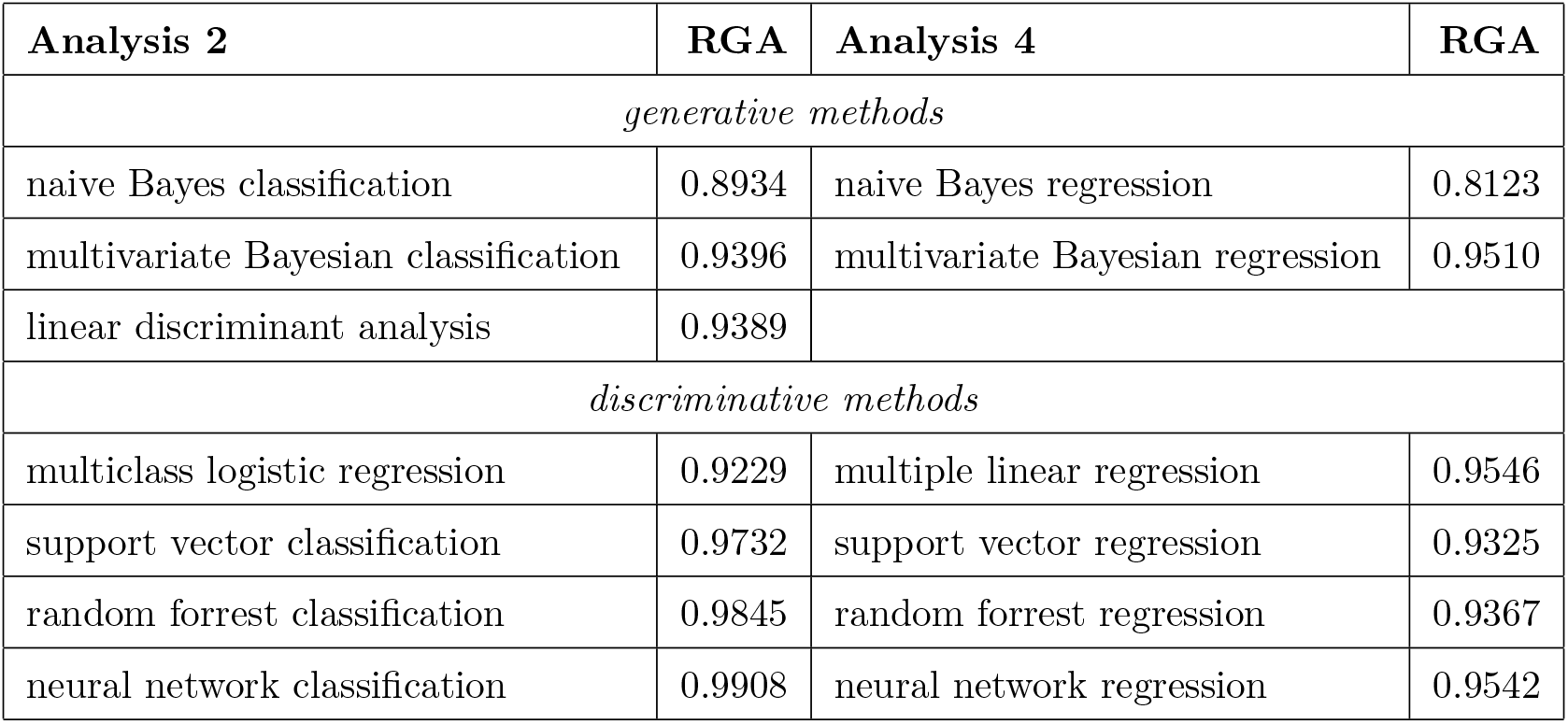
Analyses: comparisons with alternative approaches. Predictive performance for generative and discriminantive techniques when analyzing data from Analysis 2 and Analysis 4. For details regarding MBC and SVC, see Figure 8, and for details regarding MBR and SVR, see Figure 11. In all cases, predictive performance is given as rank graduation accuracy (RGA) [48].

In Analysis 4, MBR (RGA = 0.9510) and LinReg (RGA = 0.9546) perform best among generative and discriminative approaches, respectively. Except for GNB, which fails to account for covariances between features (RGA = 0.8123), differences between regression techniques are negligible, all performing slightly below the level of MBR and LinReg (see Table 4).

#### Conclusion

MBI-based predictive analysis for classification and regression reaches predictive performance comparable with that of established (generative or discriminative, Bayesian or non-Bayesian) ML techniques in empirical data sets.

## 5 Discussion

In this section, we discuss some aspects pertaining to multivariate Bayesian inversion (MBI). We categorize these aspects into “advantages”, i.e. benefits of MBI over other approaches, “disadvantages”, i.e. shortcomings relative to other approaches, and “extensions”, i.e. possibilities for increasing the applicability of MBI.

### 5.1 Advantages

#### 5.1.1 Explainability and robustness

##### → Examples: Simulations 1-4, Analyses 1-4

The proposed MBI framework aligns well with the principles of the SAFE machine learning approach [62]. In contrast to discriminative algorithms, where model weights typically lack a direct substantive interpretation, MBI is based on an explicit generative model in which parameters quantify the effect of label variables on observed features (see Section 2.1). This means that the regression coefficients can be interpreted in a straightforward manner as conditional dependencies of features on labels, thereby providing a transparent link between model structure and domain-relevant mechanisms. This property enhances *interpretability* not only at the level of individual parameters, but also at the level of the full model, facilitating scientific insight and supporting informed decision-making.

With respect to *accuracy*, MBI demonstrates competitive performance relative to widely used machine learning approaches. Empirical evaluations show that MBI either outperforms or achieves comparable results to established methods, particularly support vector machines, across multiple real-world data sets (see Sections 4.1-4.4). This indicates that the generative modeling strategy underlying MBI does not come at the cost of predictive performance. Instead, it combines strong accuracy with the additional benefit of probabilistic outputs, suggesting that high predictive quality and principled statistical modeling are not mutually exclusive, but can be jointly realized within a unified Bayesian framework.

Finally, MBI exhibits favorable properties in terms of *sustainability*, understood here as robustness and reliability of inference. By basing predictions on full posterior probabilities rather than point estimates, MBI enables balanced decision-making that naturally accounts for uncertainty. This probabilistic treatment reduces the risk of overconfident or unstable predictions and supports more robust conclusions, particularly in settings with limited or noisy data. The robustness of this approach is further corroborated by comparisons based on the robustness-focused RGA measure [48] in which MBI shows consistent and stable performance compared to alternative methods (see Sections 3.5 and 4.5). Together, these characteristics highlight MBI as a sustainable machine learning approach in the SAFE sense, combining robustness with principled uncertainty quantification.

#### 5.1.2 Classification and regression

##### → Examples: Simulations 1-4, Analyses 1-4

Unlike other approaches that are solely designed for discrete prediction, e.g. logistic regression ([12, sec. 4.3]), or continuous prediction, e.g. multiple linear regression ([12, sec. 3.1]), the MBI framework can be used for classification into discrete categories as well as for regression of continuous variables (see Figure 13); it has that in common with convolutional deep neural networks [7, 8], support vector machines [63, 64] or *k*-nearest neighbor algorithms [65, 66].

**Fig. 13.**
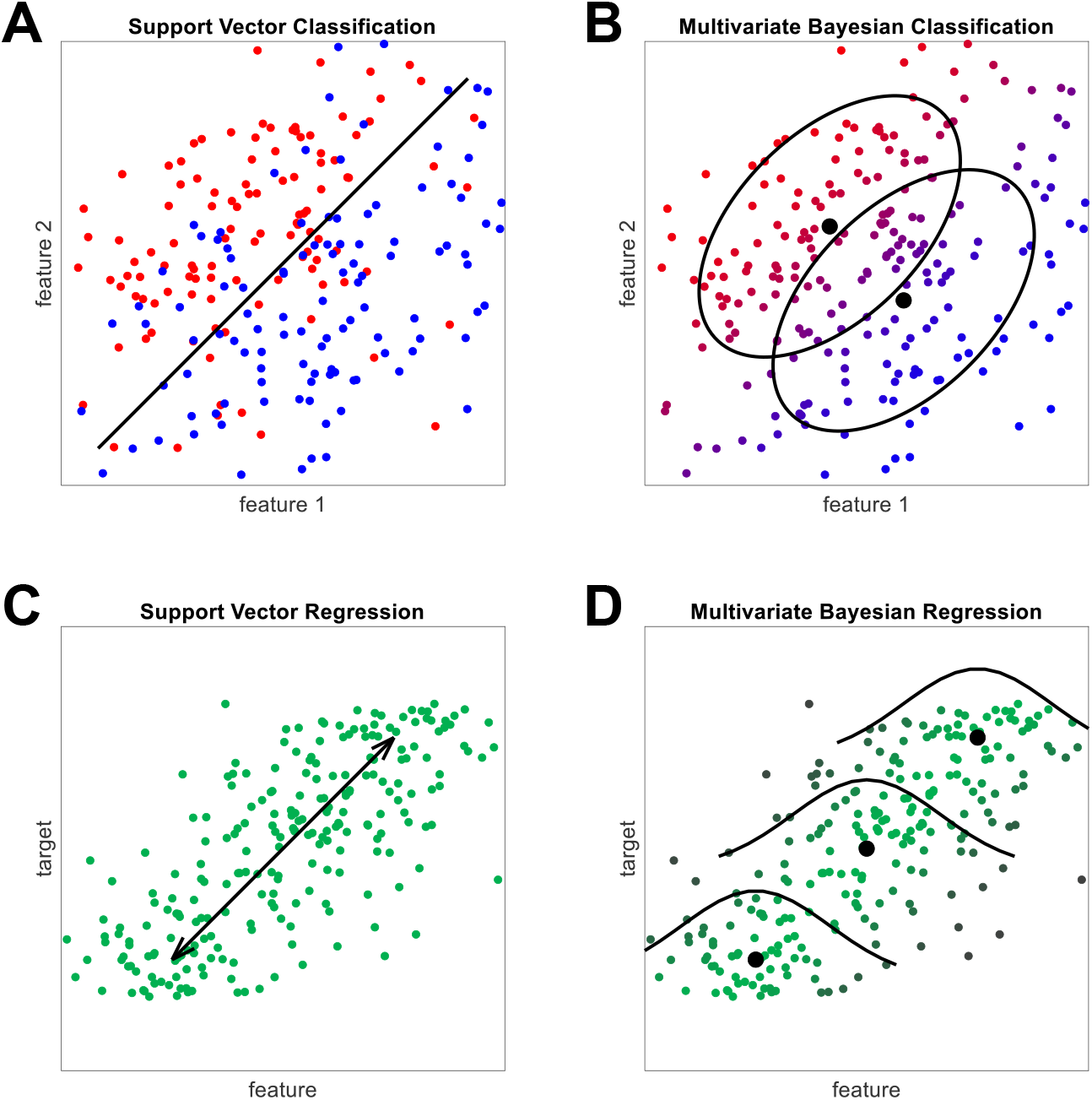
Different kinds of predictive analysis. Conceptual comparison of traditional methods (left; here: SVMs) and probabilistic methods (right; here: MBI) for classification (top) and regression (bottom), reiterating predictive algorithms vs. generative models (see Section 2.1.1). **(A)** Support vector classification searches for a hyperplane that optimally separates multivariate patterns from different classes. **(B)** Multivariate Bayesian classification estimates a probability distributions of the data, conditional on class, and uses them to calculate posterior probabilities for each class, conditional on the data. **(C)** Support vector regression searches for a projection of the data set that best matches the features onto the targets. **(D)** Multivariate Bayesian regression estimates a probability distribution of the features, conditional on the targets, and uses them to calculate posterior densities across target values, conditional on the observed features.

In multivariate Bayesian classification (MBC), the quantity of interest is treated as a discrete random variable, resulting in posterior class probabilities (cf. Figure 1B and see e.g. Figure 3A). In multivariate Bayesian regression (MBR), the quantity of interest is treated as a continuous random variable, resulting in a posterior target density (cf. Figure 2B and see e.g. Figure 6D).

Importantly, MBI works identically for both cases during the training phase, except for being based on different design matrices (see Figure 1A vs. Figure 2A). This also means that, in principle, the same model can be used for classification into categories and regression of a continuous variable at the same time, reciprocally accounting for the other, provided that both quantities of interest are included in the design matrix used in the training phase.

#### 5.1.3 Posterior probabilities

##### → Examples: Simulations 1/3/4, Analyses 2/4

Unlike many other approaches, e.g. linear discriminant analysis [41], support vector classification [5] or perceptrons [67], MBI provides posterior probabilities for the quantity to be predicted (see Figure 13); like e.g. logistic regression [68, 69] or naive Bayes classifiers [37, 70]. These are probabilities for observations belonging to particular classes in MBC and probability densities for targets having particular values in MBR.

Notably, support vector machines can be cast as probabilistic models [71]. Although probabilistic classification can be achieved using methods such as Platt scaling [72], such posterior inferences are not informed by prior distributions, but rather based on fitting a logistic regression model to the classifier’s scores [73], making it a methodologically unrelated postprocessing step not part of the algorithm itself. Moreover, the method cannot be easily extended to continuous prediction tasks for which posterior target densities from MBR are a relatively simple solution.

Probabilistic results are particularly useful for some prediction contexts, e.g. disease classification [74, 75] or individual treatment projection [76, 77]. Whereas non-probabilistic approaches for classification always make a distinct decision, a user of MBC can also refrain from making a decision, if the posterior class probability does not exceed a pre-defined threshold. Moreover, these probabilities can be combined with loss or gain functions to identify optimal decisions by application of Bayesian decision theory. Similarly, while non-probabilistic approaches for regression only give point estimates, the posterior density from MBR quantifies remaining uncertainty after seeing the data and informs the user about the precision with which the target variable is known.

#### 5.1.4 Prior knowledge

##### → Examples: Simulations 3/4, Analysis 4

The fact that MBI provides posterior probabilities comes hand in hand with the fact that it uses prior probabilities. In this way, it can integrate prior knowledge about the frequency of the distinguished classes or about the distribution of the predicted variable. For example, it might be known beforehand that one class occurs much more often than the other (e.g. healthy controls vs. patients) or that a variable tends to assume values in a specific range of its support (e.g. age or IQ).

This is especially useful in cases where it is known that the quantity of interest is constrained to a certain range of values and values outside this range are not meaningful, e.g. in brain age prediction [56]. While some approaches will inevitably also return negative values for chronological age (see [57, Fig. 2]; also see Figure 11), specification of a prior distribution can easily prevent this (see Figure 6E). In such cases, the prior distribution should be motivated from the experimental design by which the variable was controlled (see Figure 6A) or from world knowledge about the process from which the variable was sampled in case of field studies (see Figure 11).

Although this was not practiced in the present work, it is also conceivable that not only the prior model probabilities are manipulated, but the prior hyperparameters governing the MGLM parameters themselves are modified (see eqs. 9 and 17). This requires another level of prior knowledge, pertaining not just to the distribution of predicted quantities, but also to the specific mechanism connecting variables *X* and *Y*; however, it could also be helpful for regularization in the case of high-dimensional data (see Section 5.3.1).

#### 5.1.5 Feature dependencies

##### → Examples: Simulations 1/2, Analyses 1-3, Simulation 4

Like many other approaches, MBI accounts for dependencies between measured variables (between-feature covariance; Σ in eq. 3). This is in contrast to e.g. Gaussian naive Bayes [37] which assumes zero covariance between features (emulated by a diagonal matrix Σ). In principle, MBC implements a Bayes classifier for the multivariate GLM that is aware of non-diagonal feature covariance. This exploits the way how several features jointly vary as a function of class or target and enables to delineate patterns belonging to classes or targets in multi-dimensional space (see Figures 3 and 4). In fact, the between-feature covariance plays a key role, because MBI’s (unnormalized) posterior distribution is given by the ratio of two determinants (see eqs. 29 and 36), where the two matrices are hyperparameters governing the feature covariance (Ω) and the regressor covariance (Λ).

Unlike many other approaches, MBI however also accounts for dependencies between data points (between-observation covariance; *V* in eq. 3). This is important when the features to predict from are e.g. time-series data, as accounting for the correlation between data points then drastically improves prediction accuracy (cf. [16]).

#### 5.1.6 Covariate integration

##### → Examples: Simulation 2, Analyses 3/4

Many classification approaches only accept a feature matrix *Y* and a label vector *x* as inputs.^5^ This makes it difficult to handle variables which are not predictive with respect to the labels (like the features), but which might influence the features (like covariates) and thus increase predictive performance, if accounted for. MBI naturally integrates such covariates by simply adding them as regressors (see Section 2.5): For the training set, they are additional columns in the design matrix; and in the test data, they are also included as regressors, but other than the quantities of interest, their values are fixed in the model inversion process which serves to predict the quantity of interest.

We have seen that including those covariates outperforms a “prior regression” approach [45, 46] in which variation related to them is removed from the features before predictive analysis (see Figures 4 and 10). We have also seen that including the covariates outperforms employing them as features, i.e. adding them to the feature matrix (see Figure 11) in the hope that this will explain the variable to be predicted. Note that treating a variable as a covariate rather than as a feature increases the number of parameters from *r* (one weight per each covariate-as-feature-variable) to *r*× *v* (one weight per each covariate influencing each feature). This suggests that the latter approach can more flexibly account for possible effects of the covariates.

#### 5.1.7 Multi-class classification

##### → Examples: Simulation 3, Analyses 1-3

Using several examples, we have shown that MBC is capable of multi-class classification (see Figures 5 and 7). Unlike other approaches, e.g. support vector classification [78], MBI uses the same algorithm for binary and n-ary classification, i.e. a categorical design matrix (see Figure 1A and eq. 24). This means that MBC does not require separate 1-vs-1 comparisons for multi-class classification which avoids long computation times when the number of classes is very high (but see Section 5.2.5) and there is no need for a decision rule to merge several 1-vs-1 comparisons into a multi-class decision ([79]; [12, p. 339]).

### 5.2 Disadvantages

#### 5.2.1 Linear mappings

##### → Examples: Analyses 2/4

Since MBI is based on multivariate general linear models (see eq. 3), it intrinsically assumes linear relationships between labels and features, and linear effects of covariates on features. As such, it is principally restricted to such linear mappings and not able to represent non-linear relationships between labels and features in multi-dimensional space.

This is different from kernel-based methods [80], e.g. support vector machines, which also allow to exploit non-linear patterns in the data by replacing a linear kernel with e.g. radial basis function or polynomial kernels, leading to non-linear decision boundaries in cases of classification [81]. It is also a disadvantage when compared with convolutional or deep neural networks [7] which allow to represent non-linear relationships through inclusion of hidden layers or non-linear activation functions [8].

For example, it is plausible to expect that the mapping from chronological age to structural MRI data is non-linear in nature. Here, we did not explore this question, as we always applied linear SVMs as alternative methods for comparison with MBI. In an MBI-based approach, one way to faithfully capture those patterns would be to replace the MGLM-derived likelihood function by one allowing for non-linear relationships between *X* and *Y* – which would however lead to non-tractable posterior distributions and marginal likelihoods (see Section 5.3.2). Another approach would be to apply non-linear basis functions to either features or targets ([12, sec. 3.1]) or to equip MBI with a kernel-based architecture ([12, sec. 6.1]).

#### 5.2.2 Normal errors

##### → Examples: Analyses 1/2/4

In addition to linearity of the mapping from classes or targets to features, MBI assumes random variation in the features to exclusively consist of matrix-normally distributed errors. This also entails that the between-observation covariance *V* is the same across all columns and that the between-feature covariance Σ is the same across all rows of the data matrix *Y*. While the latter belief is not strictly problematic, because it corresponds to the commonly made “independent and identically distributed” (i.i.d.) assumption, the normal distribution itself may not always be a valid assumption.

The theoretical justification for normality of the errors is (i) that residual components on top of signals in the features can typically be assumed to consist of a sum of a large number of unmodeled random variables and (ii) the central limit theorem asserts that such a sum will tend to a normal distributuion ([17, sect. 3.3]). Univariate analyses depending on this justification such as multiple linear regression have turned out very robust and reliable, such that the assumption is also legitimate for multivariate approaches ([15, p. 352]). One of our examples suggests that MBI can handle moderate levels of non-normality (cf. Figure 7A). However, there will certainly exist situations in which the assumption is not warranted and in which generalized linear models may be a better choice than the MGLM [82].

#### 5.2.3 Equal covariance

##### → Examples: Simulation 3, Analyses 1-3

Through the between-feature covariance Σ, MBI can accomodate arbitrary covariance between measured signals, and because the matrix (or its inverse, *T* = Σ^−1^) is fully estimated, MBI makes no assumptions about the specific structure of this covariance. However, since the MGLM describes a stationary process, multivariate Bayesian classification assumes an identical covariance for all classes.

This means that if, for example, signals belonging to two classes are not differing in terms of their multivariate mean, but only in terms of the amount of variance or the direction of covariance between the features ([83, Fig. 3]), or if classes are differing in terms of multivariate mean *and* feature covariance (cf. Analysis 2), the estimate of the between-feature covariance will be based on the pooled data from both classes (cf. eq. 22c). Thus, MBC will likely not be able to distinguish between the classes, because the measured signals from both classes equally well fit with the pooled estimate. In each analysis, the null hypothesis should therefore precisely specify whether classes are equal only in terms of their multivariate mean (in which case such a result is statistically correct) or in terms of the distribution as such.

#### 5.2.4 Conjugate priors

##### → Examples: Simulation 4, Analyses 2-4

Multivariate Bayesian inversion uses conjugate normal-Wishart priors (cf. eq. 9) to estimate parameters of the MGLM underlying classification and regression. When a non-informative prior distribution is used (cf. eq. 17), no hyperparameters need to be specified. However, as this choice is mainly made for mathematical convenience and computational efficiency, it may also be seen as a weakness.

A fixed prior distribution reduces the model’s flexibility to accomodate uncertainty about model parameters and its capability to adapt inferences to the specifics of real-world situations. With that in mind, the potential of continuous shrinkage priors [84] or spike-and-slab priors [85, 86] might be investigated. The same holds for alternative prior distributions when infering on values of the label variable (cf. eq. 34), e.g. heavy-tailed t-distributions for regression problems [87, 88].

As these changes would make the model intractable in closed form, approximate inference techniques such as sampling inference and variational Bayes would have to be employed when using non-conjugate priors (see Section 5.3.2).

#### 5.2.5 Unimodal posterior

##### → Examples: Simulation 4, Analysis 4

When performing multivariate Bayesian regression and using a uniform prior density (see Section 3.4 and 4.4), the posterior target density typically has a global maximum and no other local maxima (see Figures 6 and 12). In other words, the posterior target density is inherently unimodal (and appears univariate normal).

This means that if, for example, the relationship between targets and features is in a way, such that two distinct values in separated regions of the target space are equally likely a posteriori, the posterior target density in MBR will most probably end up with its mode between these two values. In the worst case, this can lead to misleading conclusions, by reporting the medium value as the reconstruction, although reporting the two extreme values with equal posterior probability would have been more appropriate. Potential remedies for this are different prior densities or non-linear basis functions.

#### 5.2.6 Computational cost

##### → Examples: Simulations 3/4, Analyses 2/4

For each observation in the test set, for which a posterior class probability or the posterior target density is calculated, MBI requires the determinants of two matrices for each possible value of the predicted variable (see eqs. 29 and 36). These two matrices are the posterior precision governing the regressor covariance (Λ_2*i*_ ∈ ℝ^*p*×*p*^) and the posterior scale governing the feature covariance (Ω^_2*i*_^ ∈ ℝ^*v*×*v*^).

With the number of regressor (*p*) or the number of features (*v*) becoming very high, these matrices become very large, such that the calculation of determinants can potentially become slow. This especially holds for discretized MBR when the prior density is intended to be very fine-grained or spanning a large range of values, such that the calculation in equation (36) has to be repeated very often. Systematic comparison of computation times confirms this, showing a 5-fold or 20-fold increase of computation time for MBR, compared with SVR (see Table 5). Computation times of MBC are comparable with those of SVC and sometimes lower, especially when working with a large number of classes, e.g. in Analyses 1 and 2 (see Table 5).

**Table 5.** Computational cost. Computation times for predictive analyses based on multivariate Bayesian inversion (MBI) and support vector machines (SVM) for main analyses reported in this paper. All computations were performed using MATLAB R2025b running on a 64-bit Windows 11 PC with 32 GB RAM and an Intel Core Ultra 5 Processor 225H working at 1.70 GHz.

| Analysis | Figure | MBI | SVM |
| --- | --- | --- | --- |
| Simulation 1 | Figure 3A | 0.07 s | – |
|  | Figure 3B | 0.05 s | – |
| Simulation 2 | Figure 4C/E | 0.02 s | 0.01 s |
|  | Figure 4D/F | 0.02 s | 0.01 s |
| Simulation 3 | Figure 5B/C | 1.42 s | 0.05 s |
|  | Figure 5D/E/F | 4.23 s | – |
| Simulation 4 | Figure 6B/C | 0.29 s | 0.06 s |
| Analysis 1 | Figure 7B | 0.03 s | 0.07 s |
| Analysis 2 | Figure 8A | 01:42:33 h | 32:36 min |
|  | Figure 8B/C | 03:59 min | 06:11 min |
| Analysis 3 | Figure 10B | 0.05 s | 0.07 s |
| Analysis 4 | Figure 11, 2nd/5th column | 16.14 s | 0.94 s |
|  | Figure 11, 3rd column | 13.39 s | – |
|  | Figure 11, 4th column | 15.33 s | – |

The issue of excessively long computation times for MBR can, if necessary, be mitigated by assuming that the posterior target density is a univariate normal distribution (cf. Figures 2, 6 and 12) – in which case its parameters can be calculated from just three pairs of target value *λ* and unnormalized posterior density 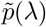.

### 5.3 Extensions

#### 5.3.1 High-dimensional data

##### → Example: Analyses 2/4

As it can be seen from the equations for posterior class probabilities (see eq. 29) and the posterior target density (see eq. 36), computation of posterior probabilities always requires the calculation of two determinants, the posterior precision Λ_2*i*_ governing the regressor covariance and the posterior scale Ω_2*i*_ governing the feature covariance. Note that, when the number of features *v* is larger than the number of observations *n*, the matrix Ω_2*i*_ will be ill-conditioned and not invertible (cf. eq. 22), such that its determinant is zero and the posterior probabilities are not defined.

One reason for this is that the prior scale matrix is set to a *v* × *v* zero matrix Ω_0_ = 0_*vv*_ (cf. eq. 17) which has the advantage that MBI does not require user-specified hyperparameters, but causes the posterior scale matrix Ω_2*i*_ to become singular. This can be avoided by setting the prior scale matrix to a scalar multiple of the identity matrix Ω_0_ = *ω*_0_ *I*_*v*_. The posterior scale matrix will then become a mixture of this prior covariance term and data-based covariance terms (cf. eqs. 11 and 22):

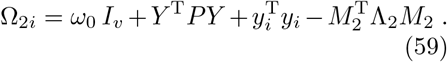

The magnitude of *ω*_0_ will determine the extent to which the features are regarded as independent signals, while the remaining terms are incorporating possible dependencies between the features based on the measured signals. Conceptually, imposing this prior knowledge on *T* via Ω_0_ is equivalent to regularizing our estimate of the between-feature covariance, similar to Tikhonov regularization in ridge regression [89, 90]. Provided that the dimensionality of Ω_2*i*_ (*v* × *v*) allows for calculation of matrix determinants in a software package implementing MBI, this can allow for prediction of classes or target variables from high-dimensional data (*v*≫ *n*), i.e. data sets with much more features than observations.

#### 5.3.2 Non-linear mappings

##### → Example: Analyses 2/4

As we have noted above (see Sections 5.2.1 and 5.2.2), MBI is restricted to linear mappings from regressors to features and matrix-normally distributed errors. Sometimes, such linear mappings are not possible or not realistic to assume, but specifying a non-linear model often renders the joint likelihood intractable and the posterior distribution cannot be expressed in closed form [12, 28, 91]. However, provided that such an alternative likelihood function *p*(*Y* |*X, θ, m*) and a prior distribution *p*(*θ* |*m*) are given mathematically, MBI can be finessed to handle non-linear mappings using one out of two strategies.

The first strategy, *sampling-based inference*, calculates the posterior distribution (in the training data) numerically and approximates the marginal likelihood (in the test data) by drawing from the prior and inserting into the likelihood [92, 93]. The second strategy, *variational Bayesian inference*, assumes the posterior distribution (in the training data) to factorize over sets of parameters [32] and calculates the variational free energy [94] as a lower bound to the marginal likelihood (in the test data).

In conclusion, while MBI does not allow for non-linear relationships between label variables and observed features, specifying an appropriate likelihood function and prior distribution allows to perform non-standard MBI with non-linear mappings [39, 95] via sampling or variational approaches.

## 6 Conclusions

We have systematically reviewed fully Bayesian inference for the multivariate general linear model (MGLM) and have shown that Bayesian MGLM estimation can enable cross-validated prediction of discrete or continuous variables. The resulting techniques of multivariate Bayesian classification (MBC) and regression (MBR) have a number of desirable properties. Further research should investigate how the proposed framework can be extended to other statistical tasks such as clustering, expectation maximization, prediction from high-dimensional data, non-linear prediction or sampling-based inference.

## 7 Statements and Declarations

### 7.2 Research Funding

This research did not receive any third-party funding.

## 7.3 Acknowledgements

The authors wish to thank Dirk Ostwald (OvGU Magdeburg) for providing helpful comments on an earlier version of this manuscript. Additionally, the authors thank the acquisitors and curators of the Egyptian skull data set [49], the MNIST data set [50], the birth weight data set [52] and the brain age competition data set [61] for providing open access to these valuable resources.

## 7.4 Data and Code

MATLAB and Python implementations of MBI for classification and regression are available from GitHub^6^. These tools allow for both, customized train-and-test analyses as well as fully automatic cross-validated prediction.

MATLAB and Python code underlying data analysis is provided in the same GitHub repository. Empirical data can be found within this repository^7^. Simulated data can be generated using the routines in this repository^8^.

## Appendix A Likelihood function of the MGLM

The model equation of the multivariate general linear model (MGLM) reads:

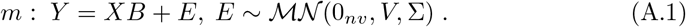

The linear transformation theorem for the matrix-normal distribution states that, if Ξ ∈ ℝ^*n*×*p*^ is matrix-normally distributed, then any (bi-)linear transformation of Ξ in the form ϒ = *A*Ξ*B* + *C* with constant *A*∈ ℝ^*r*×*n*^, *B* ∈ ℝ^*p*×*s*^ and *C* ∈ ℝ^*r*×*s*^ is also matrix-normally distributed [96, ch. 2-3]:

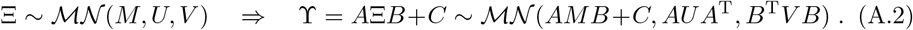

Combining (A.1) and (A.2) leads to the following distribution for *Y*

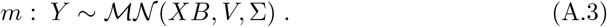

In the independent and identically distributed (i.i.d.) case, i.e. with *V* = *I*_*n*_, we can set 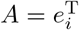, *B* = *I*_*v*_, *C* = 0_1*v*_ for *i* = 1, …, *n* and apply (A.2) to (A.3) to obtain

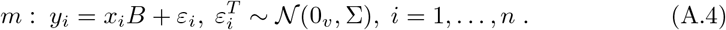

The probability density function of the matrix-normal distribution for a random matrix Ξ ∈ ℝ^*n*×*p*^ with mean *M* ∈ ℝ^*n*×*p*^ and covariances 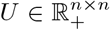 and 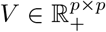 is given by [97, sect. 2.1]

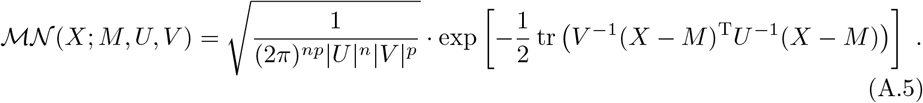

Combining (A.3) and (A.5) leads to the following likelihood function for *m*:

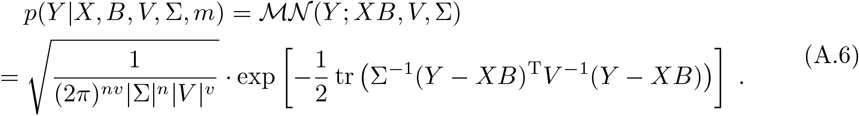

Substituting *P* = *V* ^−1^ and *T* = Σ^−1^, we finally have:

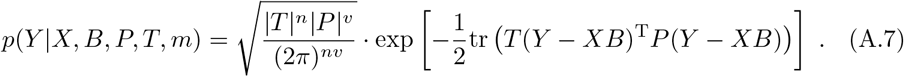

## Appendix B Posterior distribution for the MGLM

From (9), the conjugate prior for the MGLM is given by a normal-Wishart prior distribution over the model parameters *B* and *T* = Σ^−1^:

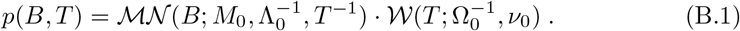

From (8), the posterior distribution is proportional to the product of likelihood function and prior density, i.e. the joint likelihood function:

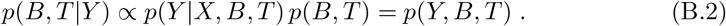

From (7), we have the likelihood function:

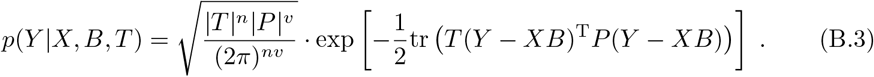

The probability density function of the matrix-normal distribution for a random matrix Ξ ∈ ℝ^*n*×*p*^ with mean *M* ∈ ℝ^*n*×*p*^ and covariances 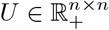 and 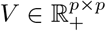 is:

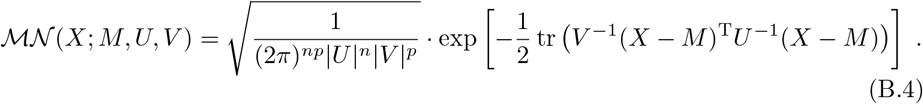

The probability density function of the Wishart distribution for a random matrix 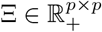 with scale matrix 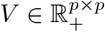 and degrees of freedom *n* ∈ ℝ, *n > p* is:

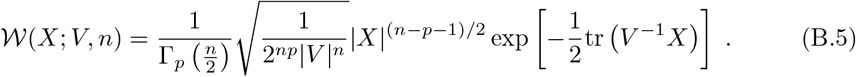

Combining (B.3) with (B.1) and plugging in (B.4) and (B.5), the joint likelihood of the model is given by:

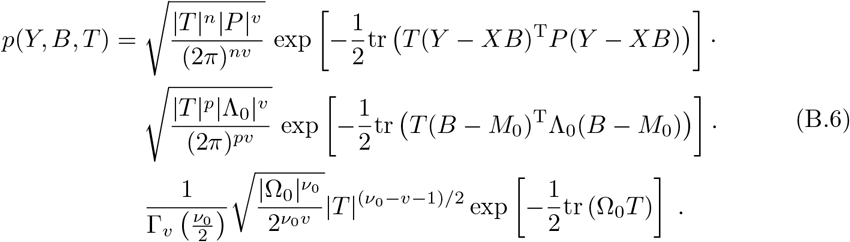

Collecting identical variables gives:

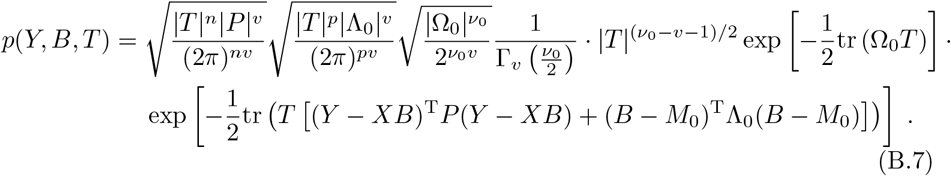

Expanding the products in the exponent gives:

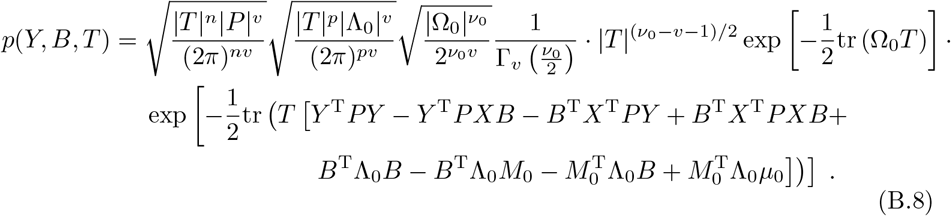

Completing the square over *B*, we finally have

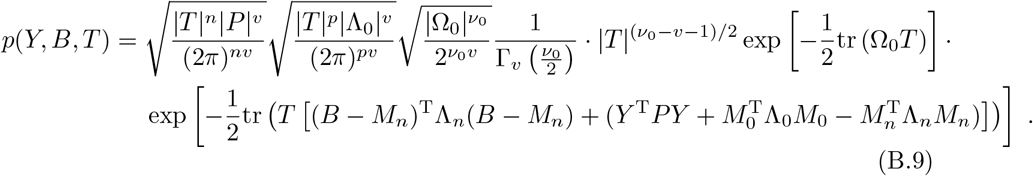

with the posterior hyperparameters

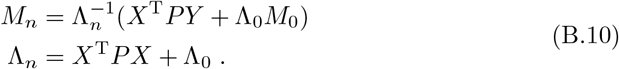

Ergo, the joint likelihood is proportional to

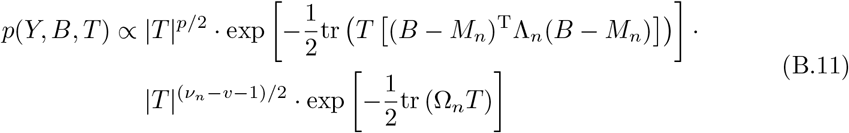

with the posterior hyperparameters

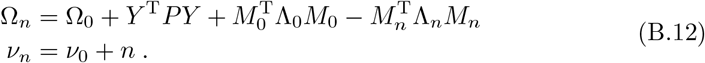

Taken together, we see that the joint likelihood (B.11) is proportional to a normal-Wishart distribution, such that the posterior distribution is also a normal-Wishart distribution^9^

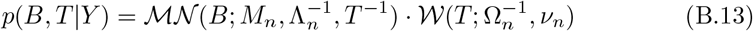

and the posterior hyperparameters are given by

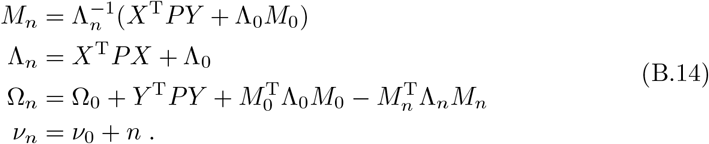

## Appendix C Marginal likelihood for the MGLM

Again, we assume a normal-Wishart distribution as conjugate prior distribution over the model parameters *B* and *T* = Σ^−1^:

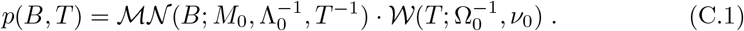

From (12), the marginal likelihood is obtained by integrating over the product of likelihood function and prior density, i.e. over the joint likelihood function:

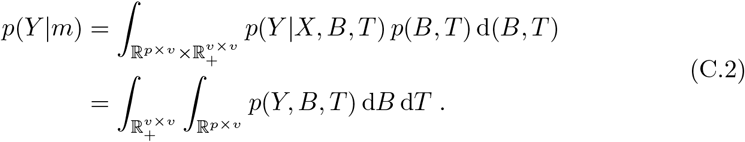

In Appendix B, the joint likelihood *p*(*Y, B, T*) has been obtained as (see eq. B.9)

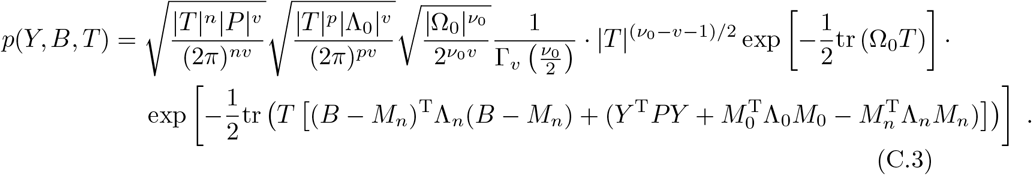

In the following, we will make use of the fact that probability density integrates to one over the whole domain of a random variable, e.g.:

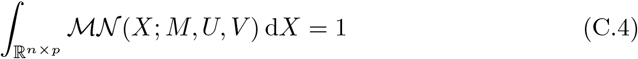

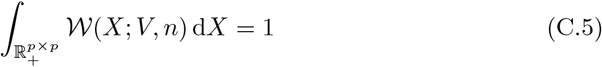

Using the matrix-normal probability density function (B.4), we can rewrite this as

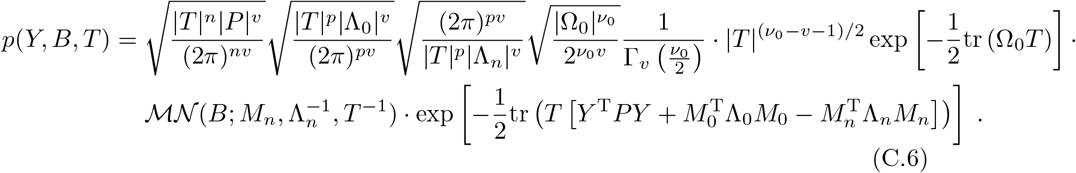

Now, *B* can be integrated out easily by appling (C.4):

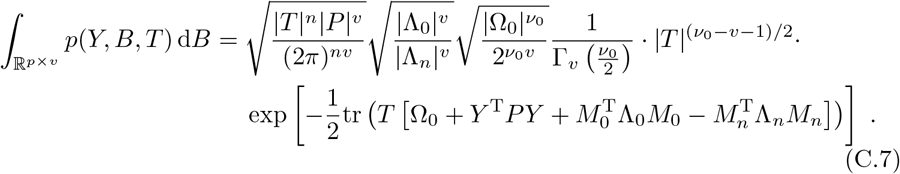

Using the Wishart probability density function (B.5), we can rewrite this as

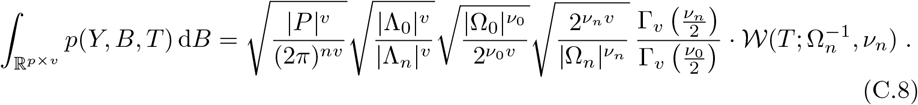

Finally, *T* can also be integrated out by appling (C.5):

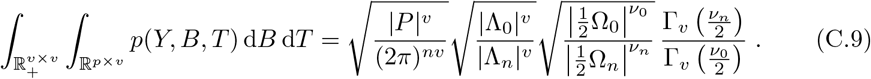

Combining (C.9) and (C.2), the marginal likelihood of the MGLM is^10^

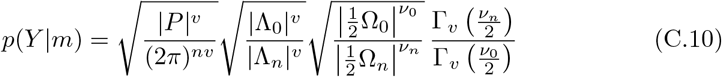

where the posterior hyperparameters *M*_*n*_, Λ_*n*_, Ω_*n*_ and *ν*_*n*_ are given by (B.14).

## Appendix D Multivariate Bayesian inversion

In this section, we use the superscript □;^(1)^ to denote quantities belonging to the training data and the superscript □;^(2)^ to denote quantities belonging to the test data.

In the first step of an MBI analysis, the data set of features and labels is

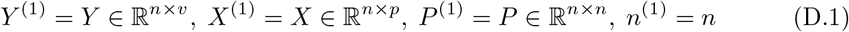

and from (17), the prior hyperparameters for the first step are given by:

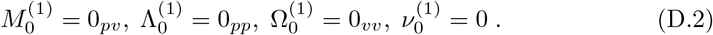

Plugging this into (B.14), the posterior hyperparameters after the first step are:

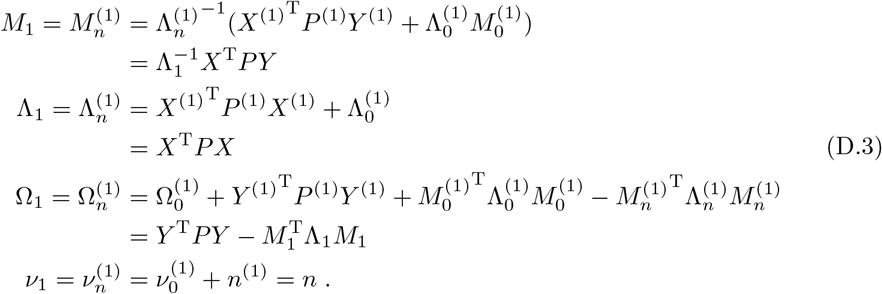

In the second step of an MBI analysis, features and labels of the single data point are

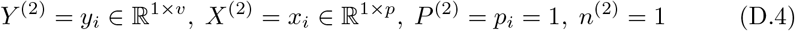

and from (D.3), the prior hyperparameters for the second step are the posterior hyperparameters obtained in the first step:

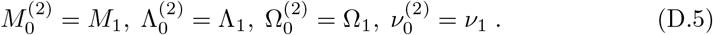

Plugging this into (B.14), the posterior hyperparameters after the second step are:

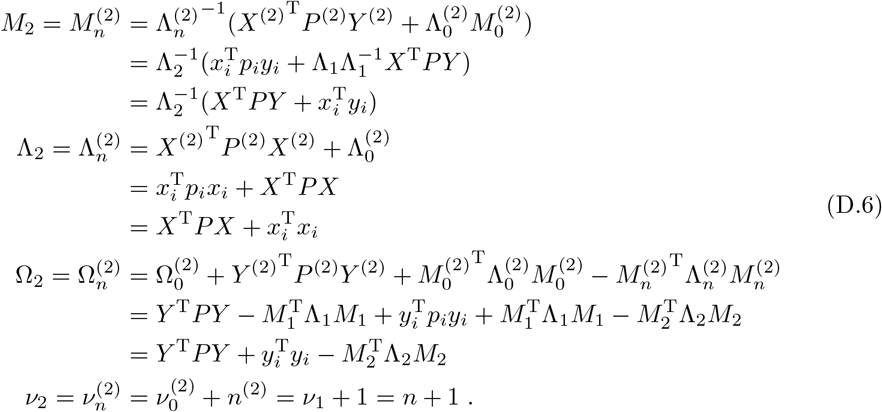

## Footnotes

1 For details, see the MATLAB implementation (https://github.com/JoramSoch/MBI/blob/main/MATLAB/ML_CV.m) or the Python implementation (https://github.com/JoramSoch/MBI/blob/main/Python/MBI.py#L341-L421) of cross-validation in the MBI GitHub repository.

2 For simplicity, this and the next matrix are specified as Toeplitz matrices with constant descending diagonals, i.e. the entries only depend on the absolute difference between row and column index (*i* − *j*).

3 The data set can be obtained from: https://github.com/JoramSoch/PAC_2019.

4 See: https://github.com/ohbm/OpenScienceRoom2019/issues/10.

5 See, for example, the Python implementation of support vector machines (https://github.com/cjlin1/libsvm/blob/master/python/libsvm/svmutil.py) or the MATLAB implementation of logistic regression (https://www.mathworks.com/help/stats/fitmnr.html).

6 See: https://github.com/JoramSoch/MBI.

7 e.g. Analysis 1: https://github.com/JoramSoch/MBI/tree/main/Examples/Analysis_1.

8 e.g. Simulation 1: https://github.com/JoramSoch/MBI/tree/main/Examples/Simulation_1.

9 The derivation of this posterior distribution is inlcuded in [98, ch. 3.4] and can also be found at: https://statproofbook.github.io/P/mblr-post.

10 The derivation of this marginal likelihood is inlcuded in [98, ch. 3.4] and can also be found at: https://statproofbook.github.io/P/mblr-lme.

